# Culture of pluripotent stem cells in microscale droplets modulates differentiation and tissue patterning towards organoids on chip

**DOI:** 10.1101/2024.10.25.620219

**Authors:** Nadia Vertti-Quintero, Clara Delahousse, Andrey Aristov, Tatiana Traboulsi, Jack-Christophe Cossec, Charles N. Baroud, Sébastien Sart

## Abstract

1

The differentiation of pluripotent stem cells (PSCs) and their self-organization into organoids are influenced by cell-cell interactions mediated by contacts and secreted molecules. These interactions are enhanced in microfluidic droplets due to confinement and small culture volumes. However, a comprehensive study on the culture of PSCs within droplets and the impact of this microenvironment has yet to be conducted. In this study, we present a droplet platform for the 3D culture of PSCs at various stages of cellular commitment. We demonstrate PSC differentiation into the three germ layers and the feasibility of organoid formation within droplets. Our findings reveal that culturing PSCs in confined volumes regulates cell fate decisions, promoting tissue patterning in gastruloids through the sequential induction of growth and migration of distinct differentiated cell populations, and facilitating the self-organization of cardiac organoids. This technological approach provides unique insights into the intrinsic factors regulating tissue self-patterning *in vitro*.

**Highlights and eTOC blurb:** *Highlights:*

- Droplet microfluidics allows expansion and supports the pluripotency of 3D aggregates of PSCs.
- Droplet microfluidics supports and regulates spontaneous differentiation into embryoid bodies.
- Droplet promotes tissue patterning in gastruloids through the sequential induction of growth and migration of mesoderm followed by ectoderm.
- Perfused microfluidic droplets support long term culture and derivation of organoids on chip. Vertti-Quintero *et al*. introduces a microfluidic droplet platform for the 3D culture of pluripotent stem cells (PSCs) at various differentiation stages. The format supports the long term 3D culture and the differentiation of PSCs -either spontaneous or directed-. This “microscale culture” can regulate PSCs fate decision, while promoting tissue pattering -as demonstrated in gastruloids polarization- and allowing self-organization towards cardioids formation.

## 3 Introduction

Organoids are small 3D structures formed from stem cells, mimicking tissues and organs with similar cellular diversity, structure, and function. Pluripotent Stem Cells (PSCs), such as induced Pluripotent Stem Cells (iPSCs) and Embryonic Stem Cells (ESCs), can self-renew and potentially differentiate into various cell types. This property enables them to serve as an abundant source of precursor cells that can form 3D organized cellular structures.

The derivation of organoids from differentiating PSCs requires them to be exposed to soluble and/or mechanical cues while being cultured within a suitable geometry to promote their organization into an organ-like 3D architecture. In an effort to reproduce such native microenvironment *in vitro*, microphysiological systems have been developed to better harness the presence of biochemical and biophysical cues that dictate PSC fate. During culture, these miniaturized systems provide, in comparison to standard methods, a better spatial and temporal control of the supply of external biochemical and biophysical cues, as well as allowing a local diffusion of cell-secreted soluble factors[1]. Moreover, microphysiological systems potentially reduce the variability of organoids production. For instance, microwells or micropatterned surfaces have been used to obtain a tighter size distribution of PSCs aggregates, favoring cells’ geometrical confinement that in turn regulate fate decision and symmetry breaking[2, 3]. Alternatively, the use of microfluidic devices has been adopted in cell culture systems, since they allow greater control of the molecular transport at the microscale[4] in comparison to systems using larger volumes. Specifically, it has been shown that the culture of PSCs in nano/micro liter scale allows a more rapid accumulation of secreted molecules around cells (*e.g*. LIF, Lefty, BMP4), regulating their processes of self-renewal and differentiation trajectories[5, 6, 7]. Moreover, by applying controlled medium perfusion at the microscale, microfluidic devices allow a tight balance between the addition of exogenous molecules and the accumulation of autocrine cues in order to direct cell differentiation while preserving their capacity for self-regulation[8, 9, 10]. This has led to the development of organoids on chip, which are usually constituted by open culture systems where the cells are physically retained using wells, pillars or hydrogel beads or capsules and continuously perfused in order to provide the cues and nutrients to cells for complete differentiation [11, 12, 13, 14, 15].

Microfluidic droplets allow the encapsulation of any kind of biological sample in aqueous droplets dispersed in oil. In particular, droplets have been widely used for cell culture in 3D format, not only to do sample partitioning and isolation, but also to get access to a new set of tools to manipulate cells and cell aggregates (*i.e*. for stimulation, sorting or observation) at the microscale[4]. In contrast to other types of micro-culture systems, the droplet format is a closed culture system that strongly favors cell-cell interactions, by promoting physical cellular contacts as well as by accelerating molecular transport[16, 17, 4]. Microfluidic droplets have been first applied for the cultivation of PSCs for controlling their encapsulation into hydrogels of different rigidities[18, 19]. In such configurations, the use of flow focusing devices allow the tight control of the gel bead size. Another strategy is the use of hydrogel capsules with liquid cores, made with triple-jet in-air microfluidics, where PSCs can be cultivated and differentiated towards organoids[15]. To note, after hydrogel gelation, PSCs encapsulated within the structures are usually cultivated in regular well plates. In this context, there is a limited number of studies that have integrated the culture and differentiation of PSCs in liquid microfluidic droplets that enable continuous tracking of PSCs evolution over time. PSCs have been cultivated into isolated or perfused liquid microfluidic droplets of volumes ranging from 200 nL to 25 *μ*l and for culture periods varying from 24 hours to 3 days[20, 21, 22, 23, 24]. These studies revealed that liquid microfluidic droplets supported the culture of PSCs at the undifferentiated state for short time, while the extent of differentiation and the efficiency of organoid formation in liquid droplets had not been investigated so far. Recently, we demonstrated that the culture of PSCs into microfluidic droplets was a mandatory step to allow the formation of gastruloids capable to give rise to embryo-like structures displaying head and trunk like structures[17, 25]. Yet, the performance of the microfluidic droplet format needed to be thoroughly characterized in comparison to standard culture conditions, in order to provide a more comprehensive understanding of the fate decisions of PSCs within microfluidic droplets.

Here, we extend the characterization of mouse embryonic stem cells (mESCs) behavior while cultivated in the droplet microfluidic platform. By benchmarking the culture of mESCs in droplets to standard 96 well plates, this study reveals that 7 *μ*l droplets are sufficient to expand and differentiate mESCs at various developmental stages. By examining key parameters such as differentiation efficiency, and organoid formation, we aim to determine how the microfluidic droplet system enhances or modifies PSC culture compared to conventional approaches. Namely, mESCs can be cultured in 3D aggregates retaining pluripotency, allowing them to early differentiate into embryoid bodies (EBs), patterning them into gastruloids and forming cardiac organoids. Altogether, the results of this study demonstrate that culturing PSCs in microfluidic droplets has the potential to regulate cell differentiation and 3D self-organization towards organoids on chip.

## 4 Results

### 4.1 A microscale droplet platform for 3D culture of Pluripotent Stem Cells

A droplet microfluidic platform dedicated to the culture of mESCs was developed. As the droplet volume had to be adapted to specific cell biochemical needs[4], we first investigated the minimal droplet volume that would allow the aggregation, and thus the formation of 3D pluripotent cellular structures of mESCs. A fixed number of cells (*i.e*. 300) was loaded within droplets of volumes ranging from 50 nl to 30 *μ*l. This was done by using our previous microfluidic platforms[26, 27, 28] to form nanoliter scale droplets (50-800 nl) and standard hanging drops for larger volumes (1-30 *μ*l). After 24 hours the drops containing cells were observed in order to assess if cells had aggregated. As it is shown in Figure 1a, for volumes below 7 *μ*l the mESCs did not manage to form aggregates. Consequently, a volume of 7 *μ*l was chosen for the culture of mESCs in 3D for the following experiments in droplets. Then, we fabricated a microfluidic platform that displayed the main features of our previous designs, namely: a large number of traps for several 3D cultures in parallel, compatibility with live imaging and allowing biological isolation and the temporal regulation of each droplet content[26, 27, 28]. To capture 7 *μ*l droplets, we designed 80 hexagonal prism traps of height (*h*) 4 mm, edge-to-edge distance (*l*) of 2 mm that are patterned on the top part of a main chamber of 1 mm height (Figure 1b, c). Of note, increasing the droplet volume, as well as the dimensions of the traps and the channels, is associated with important changes in the physical process driving droplet trapping in comparison to our previous devices[26, 27, 28]. The contribution of the buoyancy (*i.e*. due to the lower density of aqueous droplets in comparison to fluorinated oil) becomes comparable when contrasted to the capillary forces (*i.e*. due to the droplet confinement within shallow channels) for the trapping of droplets of this size in these anchors. Because of it, the droplets are first retained at the bottom part of the traps by capillarity, then they are moving to the upper part of the traps due to buoyancy. This feature was used for controlling droplet trapping and pairing in a single trap as shown in Figure 1d.

**Figure 1:**
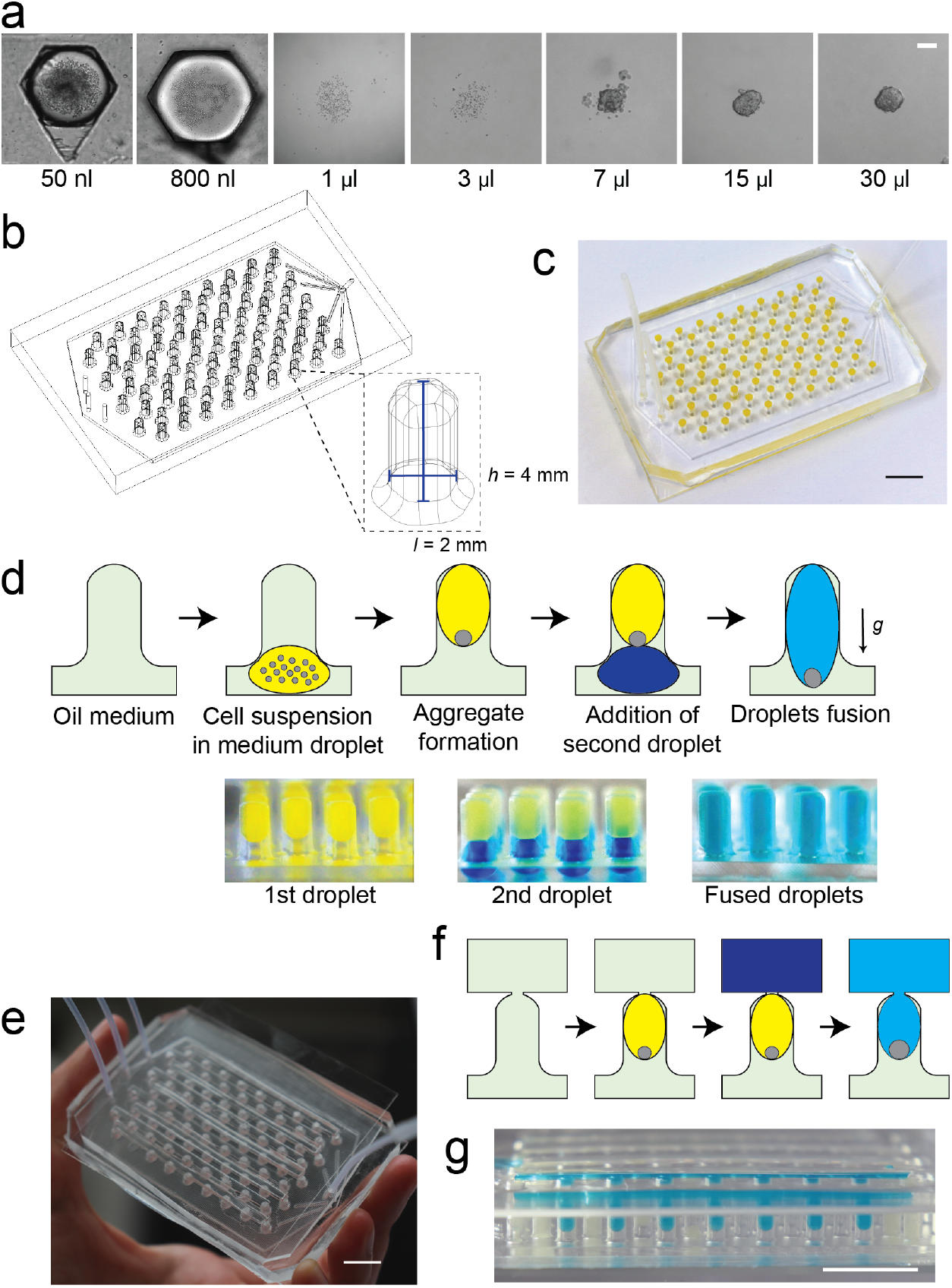
Droplet microfluidic platform for the culture of mESCs aggregates. **(a)** Bright-field images of droplets of various volumes, each containing 300 mESCs, taken after 24 hours to observe the formation of cell aggregates. Scale bar represents 50 *μ*m. **(b)** Schematic of the microfluidic device containing 80 traps, each one designed to retain a 7 *μ*l droplet. **(c)** Photograph of the microfluidic device. **(d)** Schematic of the culture of mESCs aggregate on a single microfluidic trap. First, droplets of cells suspended in culture medium are anchored in a trap. Cells settle at the bottom of the trapped droplet and form an aggregate in around 6 hours. At posterior times, a second droplet (containing *e.g*. a biomaterial, culture medium, staining solution, *etc*.) can be added on demand in the trap and fused with the initial droplet in order to tune its content. **(e)** Photograph of the microfluidic device with the appended perfusion open channels on top of the design shown in (c). **(f)** Schematic of the culture of mESCs aggregate on a single microfluidic trap with the perfusion strategy. The perfusion channel is filled with culture medium and its content is fused to the initial droplets containing cells. **(g)** Photograph of a row of traps occupied by droplets and perfused from the appended channel on the top of the device. Scale bars in this figure represent 1 cm.

The experimental protocol in the microfluidic device begins by using a cross-junction to disperse plugs of 7 *μ*l containing cells suspended in culture medium into continuous fluorinated oil containing a surfactant to stabilize the aqueous droplets [17, 25]. As the diameter of the droplets is larger than the height of microfluidic chamber, the droplets adopt a ’pancake’ shape while entering into the microfluidic device. Then, when a droplet encounters a trap, whose diameter is larger than the height of the main chamber, part of its surface energy is released when it enters partially into the trap. This process promotes the capture of the droplets in the bottom part of the traps, temporarily blocking the capture of any additional droplet. Captured droplets then move to the upper part of the trap by buoyancy. The cells in suspension can come together in the concave bottom of the hanging droplet by gravity, promoting their aggregation (See Supplementary Movie 1). Afterwards, when initial droplets are in the top of the trap, a second droplet of maximum 7 *μ*l can then be captured in each trap. By using an interface destabilizing agent (*e.g*. perfluoro-octanol, PFO), the two droplets trapped in the same anchor can be fused and their content can mix. Thus, this feature allows the temporal control over the droplet contents, as illustrated in Figure 1d. To note, in our experimental setup the intra-population variability can be assessed by comparing aggregates from different droplets within one experiment, as well as the inter-experimental variability among different chips.

Finally, isolated aqueous droplets are prone to nutrient and growth factors limitation for long periods of time. In order to extend the culture period, the microfluidic platform can be coupled with a perfusion system to replenish culture medium and/or addition of biochemical cues. The droplet perfusion on-chip is achieved by appending, on top of the traps, rectangular open channels of 2 mm width and 5 mm height along the rows of traps (see Figure 1e). Such channels are connected to the traps below them by circular orifices of 1.25 mm diameter. This configuration allows to retain the droplets in place within the traps while allowing medium exchange by diffusion from the perfusion channel, after droplet fusion (Figure 1f, g). Such open microfluidic channels[29] enable easy medium refreshment and addition of chemical cues at later time points.

### 4.2 Microscale droplets support the culture of undifferentiated mESC aggregates and regulates differentiation of embryoid bodies

After establishing the parameters for forming mESCs aggregates in droplets, we verified that this novel culture format could sustain the proliferation of pluripotent aggregates for few days. In order to ensure appropriate growth and viability of cells, aggregates cultured in droplets inside the microfluidic chip were compared to those cultured in standard round-bottom 96-well plates (in which the working culture volume was 100 *μ*l). To retain pluripotency of aggregates, mESCs were cultivated in culture medium supplemented with 2iL (see Experimental Procedures, hereafter referred to as CM^+^). The undifferentiated aggregates were found to significantly increase their size over a 4-days culture period inside the droplets, as shown in Figure 2a and in Supplementary Figure S1b. The average aggregates’ diameter started from 100 *μ*m at day 1 (D1) to reach about 220 *μ*m at day 4 of *in vitro* culture (D4), that is, more than two fold increase (Figure 2b). Note that the aggregates’ size was tightly distributed right after their formation, with a coefficient of variation (CV) of 12.4% at D1. The size distribution remained narrow during the culture, with the CV decreasing over the time to reach 8.5% at D4. The results show that 3D culture of mESCs in droplets yields homogeneous aggregates, with limited variability among droplets in the microfluidic chip. In line with the proliferation in droplets, mESCs demonstrate high percentage of viability at D4: only 5% of the cells were positive for PI (propidium iodide) staining (Figure 2c, Supplementary Figure S1b). Importantly, we found that undifferentiated aggregates cultured in 96-well plates show similar size evolution (about a two fold increase of the diameter between D1 and D4) as well as percentage of viability than mESCs in droplets, as shown in Supplementary Figure S1a, and Figure 2c.

**Figure 2:**
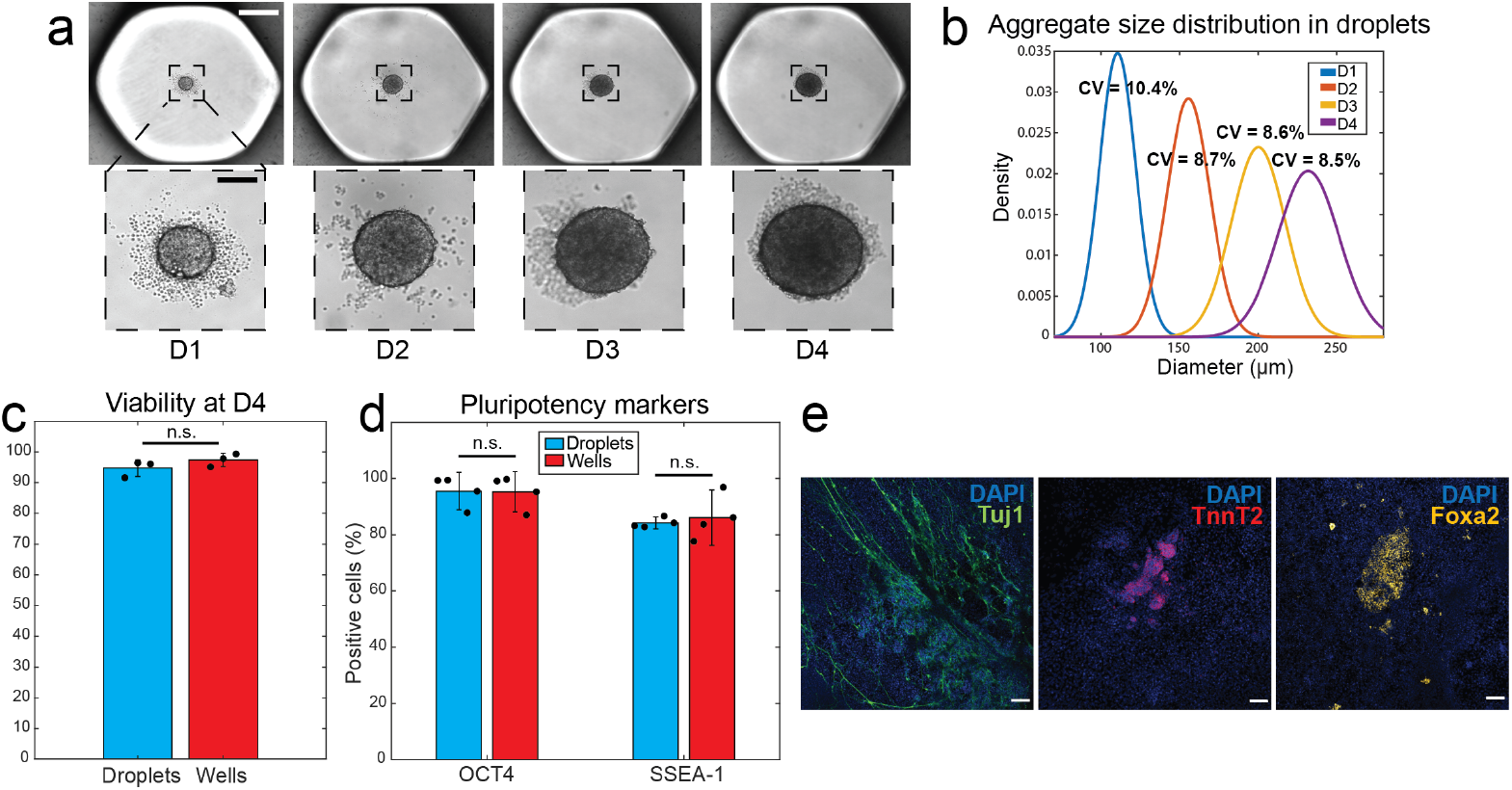
Culture of undifferentiated mESC aggregates in microscale droplets. **(a)** Bright-field images showing the size evolution of an undifferentiated mESCs aggregate cultured inside a droplet with CM^+^ over 4 days. Scale bars represent 500 and 100 *μ*m respectively. **(b)** Kernel density estimation based on the measured sizes of aggregates (*n* = 300 aggregates; N=5 chips) according to days in culture inside the chip. **(c)** Comparative of viability of cells in aggregates cultured in 96-well plate vs chip (N=3 chips; N=3 plates). **(d)** Comparison of the percentage of cells expressing Oct4 and Ssea-1 on mESCs aggregates cultured for 4 days in either chip or on a 96-well plate (N=4 chips; N=4 plates). **(e)** Expression of markers of the 3 germ layers on cells coming from aggregates in chip after being re-plated on gelatin for 8 days. Scale bars represent 100 *μ*m.

Then, the phenotype of undifferentiated aggregates was characterized by measuring the level of expression of pluripotency markers (Oct4 and Ssea-1) using imaging and flow cytometry (see Supplementary Figure S1e-f, and Experimental Procedures for details). The extent of expression of these markers was compared to mESCs cultured in 96-well plates after a 4-day culture period (Figure 2d). It was found that about 90% of the cells were positive for Oct4 and 85% for Ssea-1 both in droplets and in 96-well plates. Hence, the results demonstrate that the microfluidic platform has the capability to retain the pluripotency of mESCs similar to conventional culture systems. Moreover, to ensure that the cells retain their differentiation potential after 4 days of 3D culture in droplets, the undifferentiated aggregates were recovered and re-plated on 2D plates with culture medium without 2iL (CM^-^). This was done to test whether they had retained their capacity to produce cells from any of the three embryonic germ layers. Figure 2e shows that spontaneous differentiation of the droplet-produced aggregates yielded to the generation of Tuj1 positive cells that display neurites, typical of neural derivatives. In addition, the differentiated aggregates were composed of patches of cells expressing TnnT2 (a mesodermal markers usually expressed in developing cardiomyocytes) and Foxa2, an endoderm marker. Altogether, the droplet microfluidic platform demonstrated similar performance than 96-well plates for the expansion of undifferentiated aggregates for 4 days, albeit the culture volume was 15 times smaller.

While the droplet-format enables the culture of undifferentiated aggregates, it was not known if the microfluidic platform could support the differentiation of mESCs in 3D. Likewise, it was not known if differentiation in droplets would induce a bias in fate decision due to potential accumulation of autocrine signaling molecules in the absence of any exogenous morphogens to trigger differentiation. To investigate this, we derived embryoids bodies (EBs) in droplets, *i.e*. 3D mESCs aggregates that were allowed to spontaneously differentiate. We first attempted to cultivate mESCs using CM^-^ (see details on Experimental Procedures). However, after 2 days in culture under these conditions, the size of the EBs ceased to evolve and their viability was significantly lower at D4 in comparison to the 96-well plates (Supplementary Figure S2). To overcome such limitation, an enriched version of the culture medium (increased serum, amino acids, vitamins concentrations and improved buffering capacity, see Experimental Procedures) was used to derive EBs (hereafter referred as Diff medium) in droplets and in 96-well plates. Noteworthy, while Diff medium induces a significant increase in the EBs size, large necrotic areas were also found in the inner part of the differentiating aggregates. Yet, the size evolution and viability of the EBs was similar between droplets with Diff medium and well plates with CM^-^, as seen in Supplementary Figure S2. Consequently, to compare the differentiation of EBs in the two culture systems, we choose to cultivate EBs in droplets using Diff medium and using CM^-^ in the well plates, as aggregates had similar cell number, variability and size evolution, as shown in Figure 3a-c.

**Figure 3:**
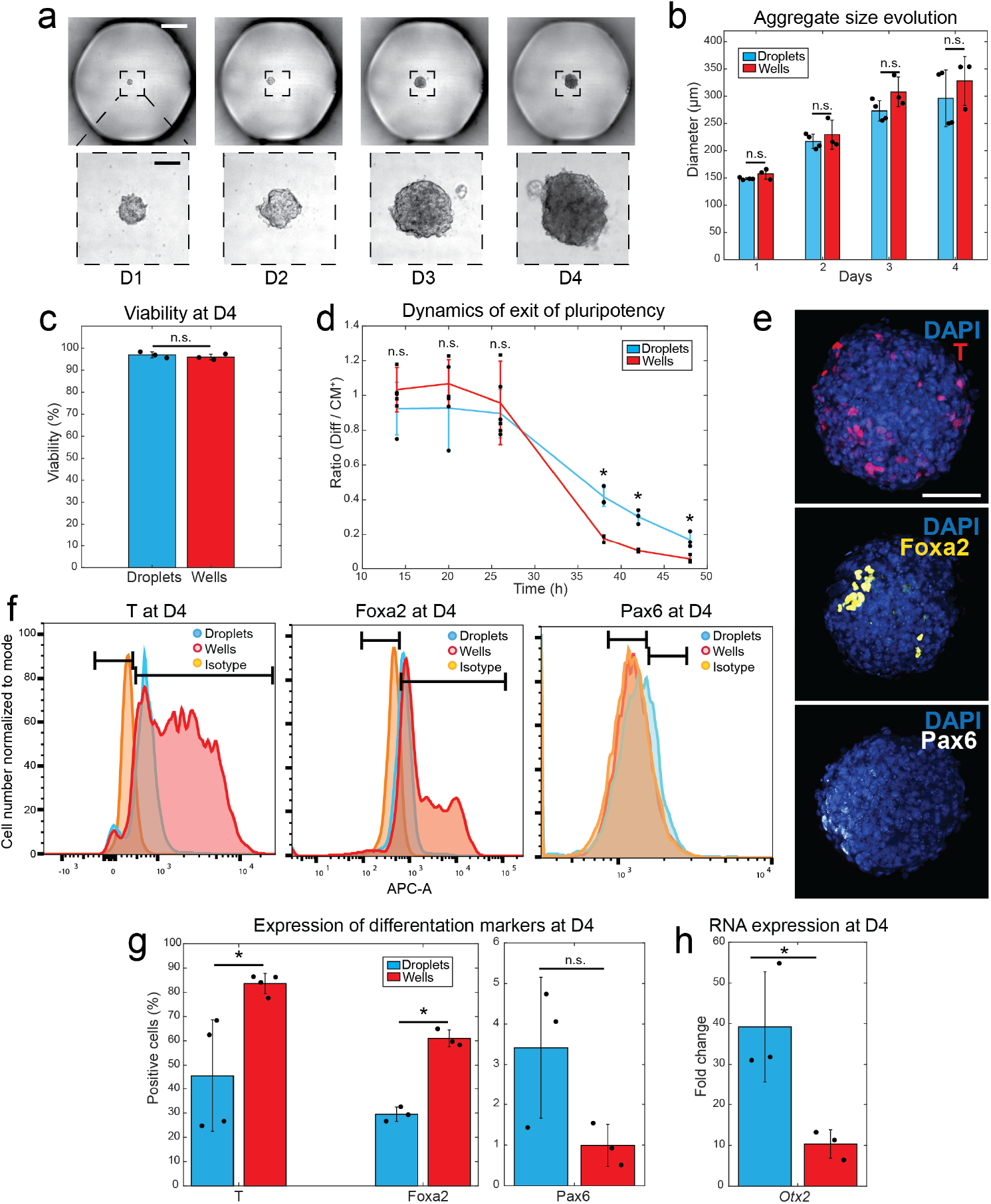
Culture of embryoid bodies in microfluidic droplets. **(a)** Bright-field images showing the size evolution of a mESCs aggregate cultured inside of a droplet in Diff medium over 4 days. Scale bars represent 500 and 100 *μ*m respectively. **(b)** Comparative of aggregate size evolution for aggregates cultured in the microfluidic chip (Diff medium) *vs* 96-well plate (CM^-^) (N=3 chips; N=3 plates). **(c)** Comparative of viability of cells in aggregates cultured in the microfluidic chip (Diff medium) *vs* 96-well plate (CM^-^)(N=3 chips; N=3 plates). **(d)** Dynamics of the expression of Rex1 in aggregates cultured in chip *vs* well plates, in (Diff or CM^-^)/CM^+^ (N=3 chips; N=3 plates). **(e)** Fluorescence images of D4 aggregates stained with markers from the 3-germ layers: Brachyury (T) for mesoderm, Foxa2 for endoderm and Pax6 for ectoderm (out of three independent experiments). Scale bar represents 100 *μ*m. **(f)** Flow cytometry plots of cells from EBs cultured in the microfluidic chip vs 96-well plate for T, Foxa2 and Pax6 (T: N=4 chips, N=4 plates; Foxa2: N=3 chips, N=3 plates; Pax6: N=3 chips, N=3 plates). **(g)** Comparative of flow cytometry results of the 3-germ layers markers expression for aggregates cultured in the microfluidic chip vs 96-well plate. **(h)** Comparison of the level of *Otx2* (ectoderm marker) expression in cells cultured either in droplets or wells (N=3 chips; N=3 plates).

We monitored the exit from pluripotency in droplets and well plates, as this would be informative of the transition from naïve to committed stages. To this end, we used a Rex1::GFPd2 reporter cell line reporter cell line to generate EBs and to form undifferentiated pluripotent aggregates as a control (see Supplementary Figure S3). The fluorescent signal reporting Rex1 (a pluripotency marker) expression was measured by live-imaging. In both culture systems, the fluorescent signal in the control experiments remained stable for 48h, consistent with the previous experiments with undifferentiated cells. In contrast, the fluorescent signal in EBs started to decrease after 24h and became almost not detectable after 48h, following a similar trend in droplets and well plates (Supplementary Figure S3, Figure 3d). Note however that the dynamics of decrease of Rex1 expression was slightly slower in the droplets than in well plates; in particular, the signal coming from GFP positive cells was about 2-3 fold brighter at 38h and 42h in the droplets than in the plates (Figure 3d).

Next, the extent of differentiation into the three germ layer lineages was measured at the protein level using immunostaining with imaging and flow cytometry (Figure 3e-g). Brachyury (T) served as a marker of the mesoderm, Pax6 of the ectoderm and Foxa2 of the endoderm, which were all previously reported to be expressed after a 4-day culture period in 96-well plates (Pax6 at a lower extend, since its expression onset was reported starting from D4[30]). Differentiated cells positive for these markers were detected after 4-days of spontaneous differentiation into the two culture systems: T and Foxa2 started to be expressed at D3, while Pax6 was only detected at D4 (See Supplementary Figure S4). Compared to 96-well plates, we quantified a drop from 80% to 40% of cells expressing T and 60% to 25% in Foxa2 in droplets, while Pax6 expression showed a trend of being higher in droplets, from 3.5% vs 1%. Noteworthy, the percentage of cells expressing T and Foxa2 in the 96 well plates followed a bimodal distribution, which suggests the presence of two distinct populations positive for these markers[31] (Figure 3f). In contrast, the percentage of T and Foxa2 expressing cells in droplets were distributed as a single peak of single positive populations. To further characterize early anterior epiblast/ectorderm differentiation[32], the level of expression of *Otx2* was compared between the droplets and the plates. A 4-fold higher expression *Otx2* expression was found in the EBs formed in microfluidic droplets in comparison to the wells (Figure 3h), which suggests increased ectoderm commitment upon confined culture.

Thus, the results demonstrated that the microfluidic droplets support the early steps of differentiation of mESCs in 3D, albeit an enriched version of the culture medium is required to sustain EB proliferation and viability. While the platform allows the exit from pluripotency in a similar manner as the 96 well plates, it regulates cell fate decision by yielding to distinct differentiation signatures in the absence of exogenous cues.

### 4.3 Culture in microscale droplets induces the formation of antero-posterior polarized gastruloids by regulating tissue growth and flow

Gastruloids are 3D cellular structures that recapitulate gastrulation, a developmental process in which the embryo’s symmetry is broken. Gastrulation gives rise to a bilateral organization composed of the primitive streak on one side and definitive ectoderm at the opposite pole. However, gastruloids generated by conventional methods (*i.e*. a pulse of chirion) usually lack anterior development due to exposure to high levels of Wnt agonists that inhibit defintive ectoderm differentiation[33]. Starting from a population of hypoSUMOylated mESCs and in the absence of any exogenous morphogen, we previously demonstrated that the droplet format promoted the self-formation of elongated gastruloids displaying similar antero-posterior polarity as a developing embryo[17]. The anterior pole of these gastruloids was formed with ectodermal cells (Pax6, Sox1 positive cells), while mesodermal cells (T positive) were localized at the posterior pole. The middle section of the aggregates was composed of Foxa2 positive cells, which formed an epithelium surrounding a lumen. Such structural organization within these type of gastruloids was not achievable using other types of culture systems such as AggreWell™, thus demonstrating the unique developmental cues provided by the culture inside droplets[17].

Here, we first verified 3D self-organization and tissue patterning of PSCs inside droplets and compared their yield efficiency of gastruloids formation to the 96-well plates. To generate gastruloids, about 120 cells derived from adherent spheroids generated from mESCs subjected to two waves of hypoSUMOylation (hereafter referred as hypoSUMOylated mESCs, see details in Experimental Procedures) were seeded in the droplets and plates for a 4-day culture period in GIM (Gastruloid Induction Medium)[17]. In these conditions, starting from the third day of culture, some aggregates broke their initial circular symmetry towards elliptical shapes, as shown in Figure 4a. Gastruloids are usually defined as 3D structures that exhibited axial elongation and a polarized expression of T[33], and we used this criteria to identify them. Using hypoSUMOylated mESCs derived from a Sox1:GFP-T:mCherry reporter line, we scored a significantly higher yield of gastruloids cultured in droplets than in 96-well plates (55% *vs* 20%, Figure 4b). As such, the results confirm that cell confinement in droplets provides a unique micro-environment to regulate 3D structural organization and fate decision within PSC aggregates.

**Figure 4:**
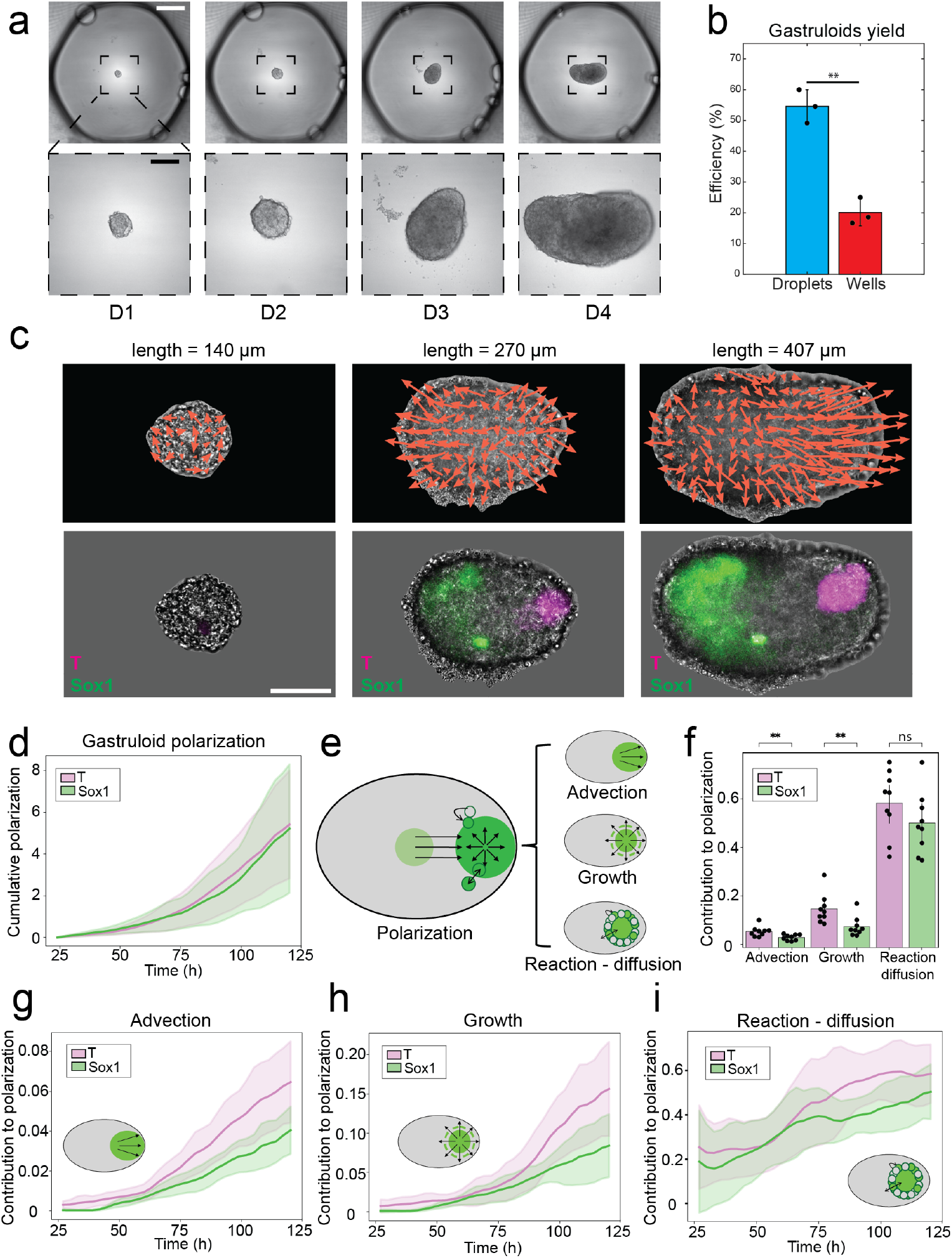
Microfluidic droplets promote gastruloid formation by regulating tissue growth and flow. **(a)** Bright-field images showing the size evolution of a hypoSUMOylated mESCs aggregate cultured inside of a droplet in GIM over 4 days. Scale bars represent 500 and 100 *μ*m respectively. **(b)** Comparative of gastruloid yield of the Sox1eGFP-TmCherry reporter cell line when cultured in droplets *vs* in wells (N=3 chips; N=3 plates). **(c)** Averaged local velocity field over gastruloids of the same length, overlaid on a bright field image of a representative gastruloid (n=9 gastruloids). **(d)** Cumulative gastruloid polarization for T and Sox1 fluorescence fields. **(e)** Schematic of the gastruloid polarization process and the model processes driving it: Advection, growth and reaction-difussion. **(f)** Endpoint measurement of the relative contribution of advection, growth and reaction diffusion. **(g-i)** Relative contribution of **(g)** advection, **(h)** growth and **(i)** reaction - diffusion to polarization in time.

Furthermore, the generation of gastruloids into stationary droplets provides a unique window to quantitatively elucidate the physical mechanisms underlying anterior-posterior patterning during gastrulation. This is due to the easy integration of culture and time lapse observations in a single microfluidic device[26]. Our protocol allows to follow the process of gastruloid formation without perturbing the culture and its monitoring, which contrasts with the “Wnt pulse” method that requires frequent medium exchanges[34]. The spatial evolution of Sox1:GFP and T:mCherry expression in the gastruloids derived from the reporter cell line was thus monitored inside the microscale droplets using live imaging for a 4-day culture period, starting from 24 to 120 hours. We measured the intensity and velocity fields of both markers, which served as a proxy to quantify the movement of ectodermal and mesodermal tissues (Figure 4c). Then, we quantified their degree of polarization (details on Experimental Procedures). The analysis demonstrated that T (T^+^) and Sox1 (Sox1^+^) positive cells separated almost symmetrically from the centroids of the gastruloids, starting from 70 hours of culture in droplets, that is, when the aggregates started to display elliptical shapes (Figure 4c).

We explored the effect of different cell-scale processes to obtain further insights on the mechanisms by which the spatial evolution of T^+^ and Sox1^+^ cells dynamically regulate gastruloid elongation. To this end, we applied the analysis proposed by Gsell *et al*.[35], in which the fluorescence intensity and velocity field measurements allow to infer the mechanisms of gastruloid polarization (P), as depicted in Figure 4d. The analysis assumes that T^+^ and Sox1^+^ cell clusters have a fluid-like behavior. By this, we mathematically decoupled the temporal evolution of the degree of polarization (*∂*_*t*_*P*) into three effects: *advection* that refers to collective differentiated cell migration; *growth* describing the proliferation of differentiated cells; and *reaction-diffusion*, which depicts the local differentiation into T^+^ and Sox1^+^ cells (see diagram in Figure 4e).

The model of gastruloid polarization into these three effects allowed us to account for most of observed cell movement over time. For instance, at 120 hours, while the induction of local differentiation had a dominant effect (55%), the proliferation contributed to 15% and the migration to about 5% of *∂*_*t*_*P* (Figure 4f). The contribution of local differentiation on *∂*_*t*_*P* was the same for both markers, while the one of advection and growth was 2-3 fold higher for T^+^ than Sox1^+^ cells. The analysis of the dynamic evolution of each effect revealed that there is a significant difference between T^+^ and Sox1^+^ cells after t = 70 hours in both advection and growth (Figure 4g,h). Indeed, the curves of advection and growth for T^+^ cells have an earlier inflection point (*τ* = 54 hours and *τ* = 38 hours respectively), thus having larger contributions to P, than those of Sox1^+^ cells (*τ* = 69 hours and *τ* = 68 hours), as shown in Supplementary Figure S5a,b. Altogether, this indicates that T^+^ cells motility and proliferation was linked with the break of circular geometry of the gastruloid and its transition towards elliptical shape (*τ* = 42 hours, as computed in Supplementary Figure S5d). Notably, at later stages of the gastruloid polarization process, we observed an increase in the rate at which the major axis of the gastruloids was extended (Supplementary Figure S5d, *τ* = 63 hours). The timing of this major axis acceleration correlates with the inflection point of advection and growth of Sox1^+^ cells (Supplementary Figure S5a,b), as well as the inflection points of both T^+^ and Sox1^+^ cells in the reaction-diffusion effect, around 60 hours (Supplementary Figure S5c).

As such, the results revealed that the culture of hypoSUMOylated ESCs into droplets promotes the formation of antero-posterior patterned and elongated structures following a biphasic temporal evolution. The first stage is determined by the motion and proliferation of T^+^ cells, which favors the formation of elliptical (polarized) aggregates. The subsequent extension of the elliptical gastruloids correlates with an increase of differentiation, migration and growth of T^+^ cells towards one pole and Sox1^+^ cells to the opposite one. Such dynamic regulation of tissue growth and flows is unique to droplets gastruloids, given the very low efficiency of gastruloid formation in 96-wells plates. The results suggest that mESCs culture in droplets provides specific cues to regulate tissue patterning within differentiating aggregates.

### 4.4 Derivation of cardiac organoids on chip within microscale droplets

The results presented in previous sections demonstrate that mESC aggregates cultivated in droplets in our microfluidic device can remain pluripotent, spontaneously differentiate into EBs, or form patterned gastruloids after a 4-day culture period. The next step was to directly differentiate mESCs into specific organoids on chip. Previously we had designed a protocol to generate cardioids from mESCs, which model heart development *in vitro*[36] that was further characterized for diversity of cardiac cells. In this protocol, which is completed after 14 days in liquid culture, ascorbic acid was used to promote cardiac differentiation. Ascorbic acid has been reported to mediate persistent ERK and FAK signaling, as well as the regulation of DNA methylation of cardiogenic-associated genes[37, 38]. As shown in Figure 5a, 3D mESC aggregates undergoing cardiac differentiation process displayed markers of cardiac progenitors from the first heart field (*Hand1*) at D5, markers of the second heart fields (*Isl1, Tbx1*) at D7, and finally *Actc1, Myl2*, both markers of differentiated cardiomyocytes, as well as endothelial market *Pecam1* and epicardial marker *Wt1* at D10, as quantified by RT-qPCR (Supplementary Figure S6). As such, the dynamics of cardiac marker expression was similar to *in vivo* embryonic heart development[39, 40]. Notably, the 3D aggregates started to spontaneously contract after 10 days in culture (about 100% of the population was beating) in 96-well plates, at a frequency of about 92 ± 24 beats/min (measured by image analysis, shown in Figure 5b, and Supplementary Movie 2). At the end of the culture period (D14), the cardioids expressed Tnnt2 and *α*-actinin (two makers of mature cardiomyocytes), CD31 (an endothelial marker) and Wt1 (an epicardial marker), as shown in Figure 5c.

**Figure 5:**
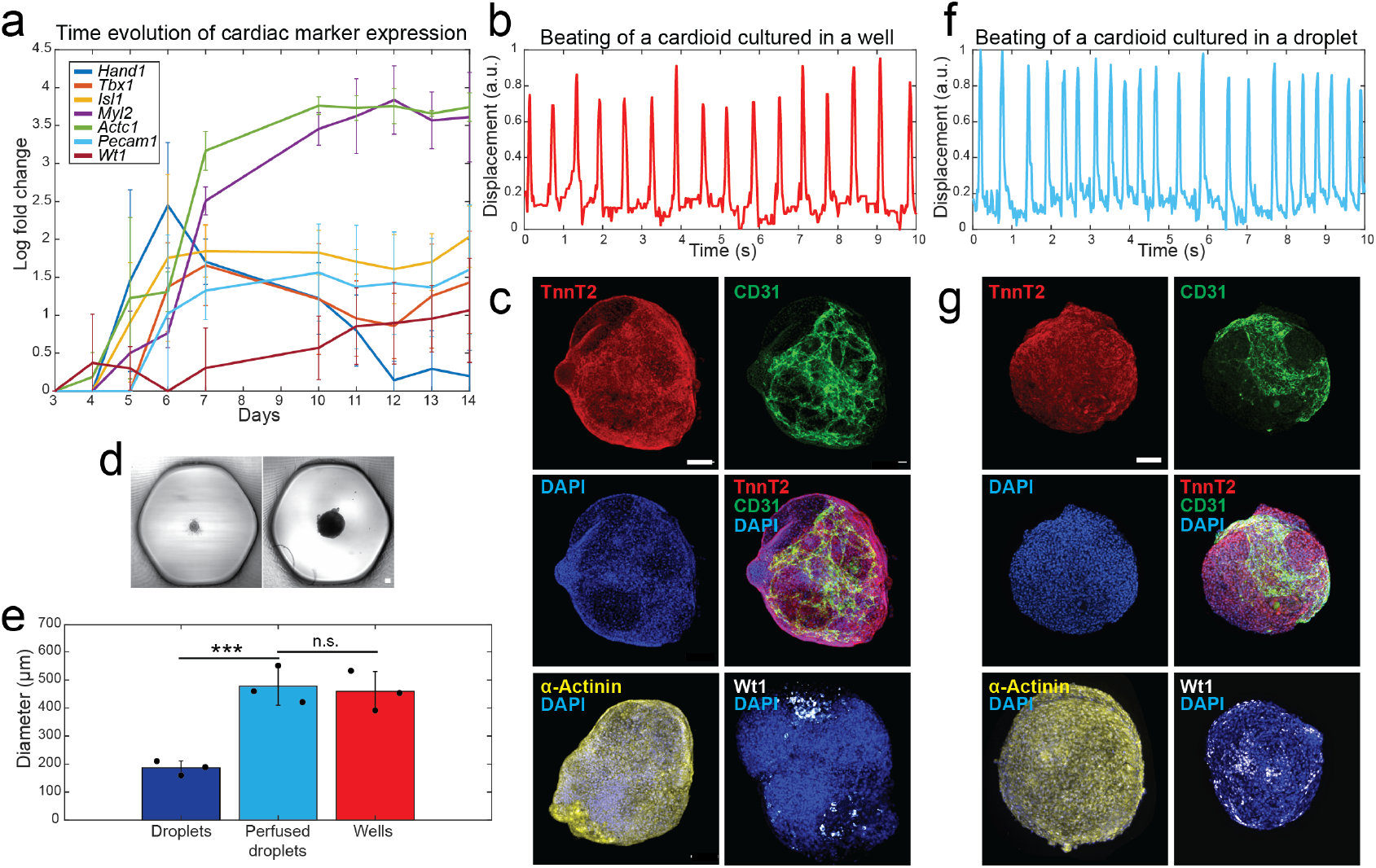
Generation of cardioids in microfluidic droplets. **(a)** Time evolution of the expression of markers for heart progenitors of the first (*Hand1*) and second heart field (*Tbx1, Isl1*), of mature cardiomyocytes (*Actc1* and *Myl2*) as well as endothelial (Pecam1), epicardial (*Wt1*) (N=3 plates) (See Supplementary Figure S6 for individual curves). **(b)** Representative displacement pattern of cardioids due to beating, from 96-well plates at day 10 of differentiation. **(c)** Fluorescence images of day 14 cardioids generated in 96-well plates and stained with markers for cardiomyocytes (TnnT2, *α*-actinin), endothelial (CD31) and epicardial (Wt1) cells (out of three independent experiments). **(d)** Representative images of D10 cardioids generated in droplets without (left) and with (right) perfusion. **(e)** Diameter of cardioids at day 10 in non-perfused and perfused droplets, as well as 96-well plates (N=3 non-perfused chips; N= 3 perfused chips; N=3 plates). **(f)** Representative beating pattern of cardioids from perfused droplets at day 10 of differentiation. **(g)** Fluorescence images of day 14 cardioids generated in perfused droplets and stained with markers for cardiomyocytes (TnnT2, *α*-actinin), endothelial (CD31) and epicardial (Wt1) cells (out of three independent experiments). Scale bars represent 100 *μ*m.

In order to demonstrate the possibility of generating organoids within the microfluidic platform, we translated the protocol for cardiac differentiation to droplets. About 300 mESCs were first seeded and cultivated for 10 days in droplets using ascorbic acid. However, when mESCs were cultured in isolated droplets, the aggregates maintained a constant size after 4 days of culture of about 200 *μ*m in diameter until day 10 (Figure 5d). In accordance, a perfusion system that would enable the renewal of droplet content over time was implemented (Figure 1e-g). By perfusing cardiogenic differentiation medium to the aggregates formerly formed and encapsulated within droplets, through the appended channel on the top of the traps, the aggregates’ growth period was extended beyond 4 days (Figure 5d). In this manner, the organoids cultivated in perfused droplets reached a diameter similar to those cultured in 96-well plates at day 10 (about 500 *μ*m, Figure 5e). At this point of differentiation, some of these large aggregates (about 35%) in droplets started to contract, at a beating frequency higher than the cardioids generated in the 96-well plates, as shown Figure 5f, *i.e*. around 110 ± 10 beats/min (See Supplementary Movie 3). Moreover, these cardioids demonstrated expression of markers for cardiomyocytes (TnnT2), endothelial (CD31) and epicardial (Wt1) cells, similar to those generated in the 96-well plates, at day 14 of differentiation (Figure 5g). Overall, the results demonstrate that equipping the microfluidic device with a perfusion system enables the long-term culture of cardioids derived from mESCs. The resulting cardioids exhibit slightly higher beating rates and display cellular diversity comparable to those cultured in 96-well plates.

## 5 Discussion

The derivation of organoids from PSCs is usually triggered by controlled addition of exogenous morphogen signals to drive differentiation in a time resolved manner. However, the addition of such extrinsic signals usually hides the self-organizing properties of PSCs, which are driven by intrinsic secreted factors and by self-tailoring immediate cellular neighboring[41]. Droplet microfluidics offers a suitable platform for 3D cell culture[4]. It has the potential to regulate cell differentiation by promoting direct cellular contact and accelerating the molecular transport of nutrients and secreted morphogens. This creates a unique opportunity to gain deeper insights into the role of intrinsic factors in regulating the self-patterning of PSC derivatives. Notably, immobilized droplets within microfluidic devices fit with the depth of field of microscopy tools, thus facilitating quantitative spatial imaging of cell fate decision down to single cell resolution in organoids[27]. Moreover, the functionality of droplet microfluidic devices can be easily extended to apply mechanical stimuli, generate molecular gradients, or multiplex culture conditions within a single device. This flexibility allows for the recreation of precise environmental niches for tissue development *in vitro*. However, the potential of PSC culture in droplets had not yet been thoroughly investigated. In this study, we developed and tested a microfluidic platform for the culture of PSCs at different stages of pluripotency or cell commitment: retaining pluripotency, undergoing spontaneous differentiation, gastruloid formation, and directed generation of organoids under exogenous morphogen-free conditions.

### 5.1 Culture and differentiation of mESCs in anchored microscale droplets

First, we found that 7 *μ*l is the minimal droplet volume that allows the aggregation of undifferentiated mESCs. This volume is 15-fold lower than the volume usually used in 96-well plates (100 *μ*l), and it is about the working volume used for cell culture in 1534 well plates (about 10 *μ*l). However, loading 1534 well plates with cell suspension requires robotic systems and it is prone to rapid evaporation of the culture medium, which can lead to important changes in the concentration of molecules in the medium. In contrast, in our microfluidic device, since the 7 *μ*l droplets are surrounded by fluorinated oil, medium evaporation is prevented for long periods.

We found that undifferentiated aggregates expand and retain expression of pluripotency markers within droplets in a similar manner as in 96-well plates, albeit the culture volume was 15 times reduced. The results suggest that 7 *μ*l droplets contain sufficient number of signaling molecules to retain undifferentiated phenotype and basic physiological cell needs. To note, it is also possible that accumulation of endogenous secreted LIF within droplets may take part in sustaining pluripotency[42, 7].

During spontaneous differentiation, mESCs transit from naive (ground) to primed state, to finally commit into differentiated phenotypes. These transitions are associated with important metabolic changes. Naive mESCs display a bivalent metabolism, in which oxidative phosphorylation (OXPHOS) contributed to 65% and glycolysis to 35% of ATP production[43]. The priming of mESCs is linked to a metabolic shift, characterized by a decrease in the reliance on OXPHOS and an increase in glycolysis for energy production, *i.e*. cells increase glucose uptake, and the medium acidifies due to lactate accumulation[43]. In this study, we found that exit from pluripotency occurred between 24 to 48 hours in droplet cultures, after which the CM^-^ medium did not sustain further EB expansion. This was most probably due to nutrient depletion and the limited capacity of CM^-^ to buffer the pH drop. In turn, we found that Diff medium extended the culture period in droplets, up to early expression of differentiation markers at D4[44].

Spontaneously differentiated EBs expressed markers of the three germ layers after 4 days in culture within droplets and 96-well plates. Primitive streak markers, such as T and Foxa2, started to be expressed at D3, while Pax6 expression emerged at D4 in both culture systems, consistent with previous studies[30]. The levels of T and Foxa2 expression in EBs were lower in percentage and magnitude when cultivated in droplets in comparison to 96-well plates. In contrast, culture of PSCs in droplets favors the induction of ectoderm markers. The expression of primitive streak markers is known to be regulated by Nodal and Wnt signaling in a time dependent manner[45, 46]. Moreover, during spontaneous differentiation, mESCs secrete large amount of growth factors and endogenous signaling molecules[47, 7], which are regulating their fate decision. In particular, Lefty, an antagonist of Nodal signaling was reported as one of the dominant factors that was up-regulated during 2D culture into microchannels[7]. As such, accumulation of Lefty in droplets may take part in the downregulation of mesendoderm and the increase of ectoderm markers expression in droplets in comparison to the well plates[48].

Together, the results demonstrate that the droplet microfluidic platform supports the 3D culture and the spontaneous differentiation of mESCs within reduced volumes of culture, while it promotes distinct regulation of molecular signaling involved in cellular specification in comparison to conventional culture vessels.

### 5.2 Derivation of organoids in anchored microscale droplets

In this study, we explored the culture and derivation of two types of organoids: gastruloids and cardioids.

The time controlled addition of exogenous morphogens (*e.g*. agonist of Wnt) or the co-culture of mESCs with trophoblast stem cells and extra-embryonic-endoderm was reported to promote tissue patterning, leading to the formation of embryonic organoids resembling gastrulating embryos (*i.e*. gastruloids)[49, 34]. We had shown that the pre-treatment of mESCs with specific inhibitors promoting hypoSUMOylation (*e.g*. ML-792 or TAK-981) induces the formation of gastruloids with antero-posterior polarity, but only if the cells are cultivated within the droplet format[17]. The regulation of gastruloids fate decision within droplets may be a consequence of geometric confinement[50], regulation of molecular transport[9] or cell interactions with the oil-medium interface[51].

In the context of gastruloids, the culture in droplets not only allows for the protocol to be carried out, but also it enables the observation the whole process of gastruloid formation, up to five days. In this manner, we investigated the physical mechanisms by which aggregates self-organized into patterned structures within droplets. We found that the formation of gastruloids in our culture configuration is a two stages process, which results of the sequential induction of the growth and migration of the primitive streak followed by ectodermal cells. The treatment of the information from the reporter cell line Sox1:GFP-T:mCherry as fluid flows can explain the majority their gastruloids’ polarization process. Other cell types within gastruloids can also play a role in the regulation of polarization, such as endoderm cells[52] or ESC-like cells, which are source BMP4 that triggers T expression[17]. In other studies, similar results were obtained using gastruloids generated with a pulse of Wnt agonist, which demonstrated that advection of T^+^ cells accounted 30% of the initial polarization. However, the evolution of gastruloids was monitored over a much shorter time period due to the technical constraints imposed by the need of frequent medium change[35]. Here we revealed the role of droplets to regulate the proliferation and migration of Sox1^+^ cells that triggers the extension of patterned gastruloids, which can not be addressed using the chiron pulse protocol or using regular 96 well plates.

We had characterized the behavior of mESCs into confined droplets for short term culture (4-5 days). However, the generation of organoids requires longer time in culture, which largely exceeds the limits imposed by the liquid droplets format. Moreover, the protocols for generation of organoids requires frequent medium exchanges, which is not compatible with droplets, even when using droplet fusion. Indeed, while droplet pairing allows the addition of molecules in a time controlled manner[28], it does not allow the withdrawal of metabolic by-products, which would ultimately lead to pH drop and increased osmotic pressure in the medium, both known to initiate the processes of apoptosis and necrosis[4]. In order to extend the culture time on chip, there was a need to allow medium exchange within immobilized droplets. Here, this was performed by equipping the top of the chip with a perfusion channel, similar to perfused hanging drops[53]. In this study, the perfused droplets were used for the generation of cardioids, which model heart development. We used our previously published protocol[36] to generate and further characterize cardiac organoids from mESCs that demonstrate similar cellular diversity (*i.e*. cardiomyoctes, endothelial cells, epicardial cells), structural organization (*e.g*. presence of cardiac cavities) and functions (*e.g*. regular beating pattern), just as reported by other groups with hiPSCs[54, 55]. Then, this protocol was translated to the droplet format. We demonstrated that cardioids with phenotypic and functional features comparable to those generated in 96-well plates can be successfully obtained. Of note, the beating frequency of the cardioids generated in droplets is slightly higher than those generated in wells. To overcome the reduced yield of cardioids generation in droplets, future work is needed to improve mesoderm induction towards cardiac differentiation in this culture format. The regulation of Wnt signaling was demonstrated to play a major role in the generation human cardioids[55]. As such, the addition of Wnt agonists (such as chirion) at early stages of differentiation may improve the efficiency of cardiogenic differentiation in droplets.

In conclusion, this study demonstrates that droplet microfluidics supports the long-term, 3D culture and differentiation of pluripotent stem cells. Cell fate decisions that allow for tissue patterning and self-organization within organoids can be regulated in liquid droplets. By clarifying the specific cues offered by the droplet format and integrating them with time-controlled perfusion, this organoid-on-chip approach could enable the consistent development of complex, well-organized tissue-mimetic structures, with potential applications in drug screening and disease modeling.

## 6 Experimental Procedures

### 6.1 Cell culture

A murine ES-D3 or ES-R1 line (American Type Culture Collection, CRL-1934 and SCRC-1011) was maintained on 0.1% gelatin-coated 25 cm^2^ flasks (Corning) in a standard 37°C, 5% CO2 incubator (Thermo Scientific). Their maintenance medium consists of the Glasgow’s modified Eagle’s medium (GMEM, Invitrogen) supplemented with 10% ESC-screened fetal bovine serum (FBS, Invitrogen), 1 mM sodium pyruvate, 0.1 mM *β*-mercaptoethanol, penicillin (100 U/ml), streptomycin (100 *μ*g/ml) (all from Invitrogen), supplemented with 2000 U/ml LIF (Millipore), 3 *μ*M CHIR99021 (Sigma) and 1 *μ*M PD0325901 (Sigma). In the text, this medium is referred as CM^+^. CM^-^ refers to CM^+^ without LIF, CHIR99021 and PD0325901. Prior to experiments, the cells were seeded at 2-4×10^4^ cells/cm^2^ and subcultured every 2–3 days.

### 6.2 Formation of undifferentiated aggregates, embryoid bodies, gastruloids and cardioids

To generate undifferentiated aggregates and embryoid bodies, mESCs were first dissociated with TryplE (Invitrogen) in order to obtain single cell suspension. About 300 mESCs in CM^+^ (for pluripotent aggregates) or CM^-^ */* Diff (for EBs) were loaded in each droplet or seeded in each well of a 96-well (round bottom) plate treated for ultra-low adhesion (Corning) and cultivated for 4 days. Diff medium was composed of Iscove’s Derived Eagle’s medium (IMDM, Invitrogen) supplemented with 20% ESC-screened fetal bovine serum (FBS, Invitrogen), 1 mM sodium pyruvate, 0.1 mM *β*-mercaptoethanol, penicillin (100 U/ml), streptomycin (100 *μ*g/ml) (all from Invitrogen).

A mouse Rex1::GFP ESC reporter cell[44] was maintained in NDiff^®^ 227 (Takara Bio) supplemented with 1000 U/ml LIF (Millipore), 3 *μ*M CHIR99021 (Sigma) and 1 *μ*M PD0325901 (Sigma), and subcultured as above.

The gastruloids derived from hypoSUMOylated mESCs was performed as previously reported[17, 25]. Briefly, ES-R1 cells or Sox1eGFP–TmCherry double reporter derived from CGR8 ESCs[56] (refered as Sox1-T) were dissociated with Trypsin-EDTA and plated at a density of 2.5 × 10^6^) cells in 100 mm dish, using serum with LIF medium (DMEM + 15% FBS + 1000 U/ml LIF) supplemented with 2.5 *μ*M ML-792 (CliniSciences, HY-108702). Medium was replaced the next day to refresh treatment. After 48 h of treatment, plates were rinsed twice with PBS then cells were allowed to recover in serum + LIF. 5 days after the end of the first round, cells were similarly counted and plated for a second wave of ML-792. After 48 h of treatment, plates were rinsed twice with PBS then cells were allowed to recover in N2B27 + LIF medium (referred as Gastruloid Induction Medium, GIM). 6 to 8 days later, cells form three-dimensional adherent spheroids. About 120 cells derived from were loaded in 96 wells plates and cultivated for 4 days in N2B27 containing LIF[25].

To generate cardiac organoids, about 300 cells were seeded using CM^-^ in the wells of 96 well plates treated for ultra-low adhesion (Corning). After three days in culture the medium was supplemented with 100 *μ*M 2-phospho-L-ascorbic acidic (Sigma), which medium was refreshed every two days until day 14[36].

### 6.3 Microfabrication

The designs for the molds to fabricate the microfluidic chips were created using Fusion360 (Autodesk). Such molds were 3D printed in a SLA 3D printer (Form3, Formlabs), using a ClearV4 resin (Formlabs). The molds were replicated in PDMS (SYLGARD®, Dow), at 1:10 base and a curing agent ratio, cured at 65°C for at least 3 hours. After curing, the PDMS was separated from the molds, outlets were cut out and then oxygen plasma treated (Cute, Femto Science Inc.) for bonding. The PDMS slab was then bonded a 75 × 50 mm glass slide (Corning #2947) and placed in an oven set at 80°C for at least 1 hour. Finally, the chips were rendered fluorophilic by treating them with Novec™1720 (3M) and heating them at 110°C for one hour. In order to extend the culture of cells within droplets towards organoids formation, a second mold was similarly created for the perfusion channels, which would be appended on top of the initial microfluidic device. The appended perfusion channels were fabricated by casting and curing PDMS onto 3D printed molds, as above. The resulting perfusion channels were 4 cm in length, 1 mm wide and approximately 5 mm in height. The top part of the traps on the initial PDMS slab were opened using a 1.22 mm punch (WPI). The top part of the initial chip and the appended channels were plasma bounded as aforementioned. At the beginning of the experiment (during droplet loading), in order to prevent oil leaking from the perfusion channel, the PDMS scraps generated during the fabrication of the perfusion channels were temporarily placed and secured on the top of the channels to close them, then after removed to open those channels and allow medium perfusion to the droplets.

### 6.4 Cell loading within microscale droplets

An Upchurch cross-junction (PEEK, low pressure, 1/16 compression size) was used to form 7 *μ*l plugs flowed at 1,000 *μ*l/min using syringe pumps (neMESYS, Cetoni) and each containing approximately 300 (120 only for the gastruloids assay) pluripotent stem cells (PSCs) in their culture medium. The aqueous plugs were separated by 6 *μ*l plugs of fluorinated oil (FC-40, 3M) containing a fluorogenic surfactant (RAN biotechnologies) at a concentration of 0.5% v/v, and were also flowed at 1,000 *μ*l/min. The chips were placed at an angle of 45° from the horizontal to allow gravity to act as a driving force for droplet motion. The drops were then spontaneously captured in the bottom part of the traps, thus preventing other drops from being anchored in the same traps at this stage. The drops then spontaneously moved by gravity to the top part of the traps over 3-5 minutes, leaving empty the bottom part of the traps. The chips were then placed in a humidified incubator (37°C, 5% CO2) to allow PSC culture for long time periods. The performance of the droplet-microfluidic platform to promote PSC expansion, maintain expression of pluripotency markers, promote spontaneous differentiation or to induce gastruloid formation was compared to standard round bottom 96-well plates.

### 6.5 Live-dead assay and immunostaining

To quantify cell viability on chip, the droplets containing the aggregates or EBs were fused with other drops containing 4 *μ*M propidium iodide (IP) (Sigma-Aldrich) and ReadyProbes™ blue (Thermo-Fisher). The staining solution was loaded in 7 *μ*L droplets that were trapped at the bottom part of the traps. The fusion of the two droplets resulted in the dilution of the staining solution, that was composed of 2 *μ*M PI and ReadyProbes™ blue. The 3D cellular structures were incubated in the droplets with the staining dyes for 4 hours, then imaged using a motorized microscope. An equivalent protocol was applied for aggregates and EBs in 96-well plates, where 100 *μ*l of staining solution was added to the culture medium, resulting to its dilution to the half.

For immunostaining, the aggregates, EBs and organoids were first recovered from the chips and the plates, then fixed with 4% paraformaldehyde for 2 hours at 4°C. The structures were then incubated overnight at 4°C in PBS-FT (5% FBS and 0.5% Triton-X100 in PBS).

The primary antibodies (Table 1) were diluted at 1:100 in PBS-FT and incubated with the samples overnight at 4°C on an orbital rocker. After 3 washes with PBS-FT, the samples were incubated overnight with a solution of 1:100 diluted secondary conjugated antibodies (Table 1) containing 0.2 mM DAPI (ThermoFisher Scientific #R37606) at 4°C on an orbital rocker. The specificity of the primary antibodies was verified by incubating the samples with the secondary antibody alone. Under these conditions, an absence of fluorescent signal validated the specificity of the primary antibodies.

**Table 1:**
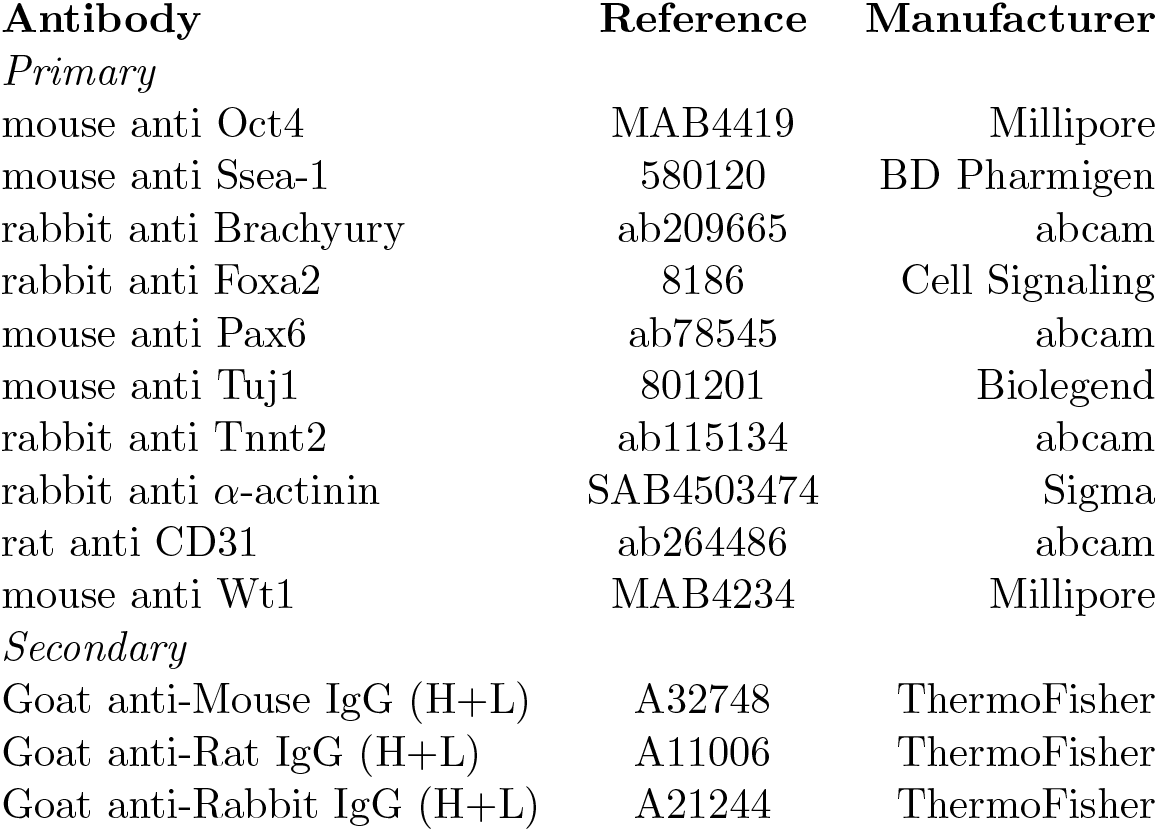
Antibodies used for immunostaining.

### 6.6 Microscopy and image analysis

The images were acquired using a motorized microscope (Ti or Ti 2, Eclipse, Nikon), equipped with a CMOS camera (ORCA-Flash4.0, Hamamatsu). Bright field and fluorescent widefield imaging were performed by illuminating the samples with a fluorescence light-emitting diode source (Spectra X, Lumencor), while for spinning disc confocal imaging the samples were illuminated with lasers (W1, Yokogawa). The images were taken with a 10× objective with a 4-mm working distance (extra-long working distance) and a 0.45 numerical aperture (NA) (Plan Apo *λ*, Nikon). Z-sections were taken every 2 *μ*m.

The images were quantitatively analyzed using Fiji or Python. Briefly, the diameter of cell aggregates and EBs was measured from microscopic images by first segmenting the images and generating ’masks’ for the aggregates. Aggregate contours have been segmented using napari-segment plugin with the gradient detector on the 4x-binned brightfield image. Centroid and major axis orientation have been extracted to cancel rotation and recenter the gastruloids. Where needed, the images have been flipped 180 degrees manually using Fiji or Python scripts and pixel-precise manual alignment performed prior to PIV analysis. Aggregates’ diameters were calculated by fitting the area of the masks to circles and resolving:

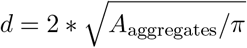

The viability of the aggregates and EBs was calculated by measuring the area occupied by PI positive cells (*A*_PI+_), which was divided by the total area of the 3D structures stained with ReadyProbes™ blue (*A*_aggregates_),

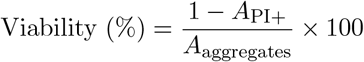

To measure the level of Rex1 expression withing aggregates and EBs, the 3D structures were first segmented from brightfield images. Then, the average fluorescent signal of Rex1::GFP reporter within each masks derived from the segmented EBs was quantified and normalized to the fluorescent intensity value at t=15 hours, using Python.

The beating of cardioids was measured from bright field movies, which were taken for duration of a 10 seconds and at frequency of 50 fps. The region of interest was then resliced, and from it, the aggregate displacement was measured using Fiji.

### 6.7 PIV and gastruloid polarization analysis

To capture the temporal evolution fluorescence fields, live images were acquired using wide-field fluorescent imaging on a Muvicyte (PerkinElmer) equipped with a 10x objectivewith a 10-mm working distance and 0.30 NA (UPlanFL N, Olympus). The microscope and sample were placed in a humidified incubator at 37°c with 5% *CO*_2_ and images were taken every three hours for 120 hours. The images were later segmented and aligned along the major axis to minimize grastruloid mouvements. Out of 11 gastruloids that were imaged, 9 were further analyzed while 2 were discarded due to miss alignment and out of frame images. The polarization analysis was conducted using the model developed in [35]. In this model, polarization is defined as the instantaneous dipole moment of either T or Sox1 fluorescence fields:

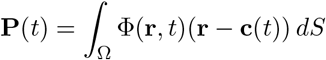

Knowing the velocity field of the cells inside the grastruloid, the temporal derivative of the polarization can be decomposed as the sum of an advection, a growth and a reaction diffusion term:

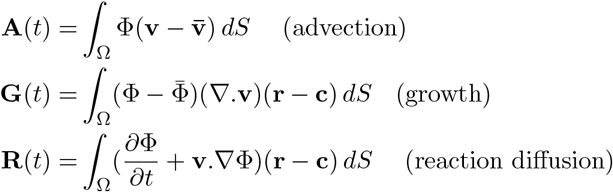

where Φ is the T or Sox1 fluorescence field, **c** is the gastruloid’s centroid and **v** is the local velocity field inside the gastruloid. PIV analysis was conducted on bright field re-aligned images of the gastruloids using PIVLab to obtain the velocity field.

For each gastruloid, the cumulative quantities of 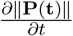, (also noted *∂*_*t*_*P*) ∥**A**(*t*)∥, ∥**G**(*t*)∥ and ∥**R**(*t*)∥ are computed. To assess the relative contributions of the advection growth and reaction diffusion terms, each of these cumulative quantities was normalized by ∥**P**(**t**)−**P**(**t**_**0**_)∥.

For the advection and growth contributions, the mean over 9 gastruloids was fitted with an exponential fit 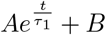. Respectively the reaction-diffusion contribution was fitted with a logistic fit 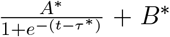, with *A* and *B*, respectively *A*^*^ and *B*^*^ two constants and *τ*, respectively *τ* ^*^referred as the characteristic time.

To measure the level of eccentricity of gastruloids, the 3D structures were first segmented from brightfield images. The aggregates were then centered and aligned along their major axis, which enabled to measure their major (a) and minor (b) axis length, using Python. The eccentricity, was calculated as follows:

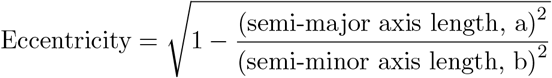

### 6.8 Flow cytometry

The aggregates and EBs were collected from the chips and plates, and then dissociated to single cells by using TriplE. The samples were then fixed using a 4% v/v PFA solution for 30 minutes and permeabilized for 10 min using a 0.5% Triton-X100 in PBS solution. The samples were then blocked with a 5% FBS (for T and Foxa2 staining) or 10% mouse serum (for Pax6 staining) diluted in PBS. The cells were then incubated with a primary antibody 1:100 diluted solution in PBS-FT solution for 2 hours, then with solution of secondary antibody at the same dilution. After washing with 5% FBS, the percentages of T-, Foxa2- and Pax6-positive cells were analyzed using a FACS LSRFortessa (BD Biosciences, San Jose, CA). To validate the specificity of the antibody staining, the distributions of fluorescently labeled cells were compared to cells stained with isotype controls.

### 6.9 RT-qPCR analysis

For the characterization of the EBs in droplets, the whole chip content or the aggregates generated from 96 well plates were collected at the end of the differentiation protocol, while undifferentiated cells cultivated in 2D using 2iL medium was used as a control. For each time point of the cardioids generation protocols, 24 aggregates were first collected from the plates. Total RNAs were extracted and purified using the RNeasy micro kit (Quiagen). 100 ng of RNA were reverse transcripted using a Quantitect Reverse Transcription kit (Quiagen). The cDNAs were amplified using a FastStart Universal SYBR Green Master Mix (containing Rox) (Roche) and primers (Eurofins Scientific, France) at the specified melting temperature (Tm) (Table 2) using a QuantStudio 3 (Thermo Fisher Scientific) thermocycler. As a negative control, water and total RNA served as template for PCR. To validate the specificity of the PCR, the amplicons were analyzed by dissociation curve.

**Table 2:**
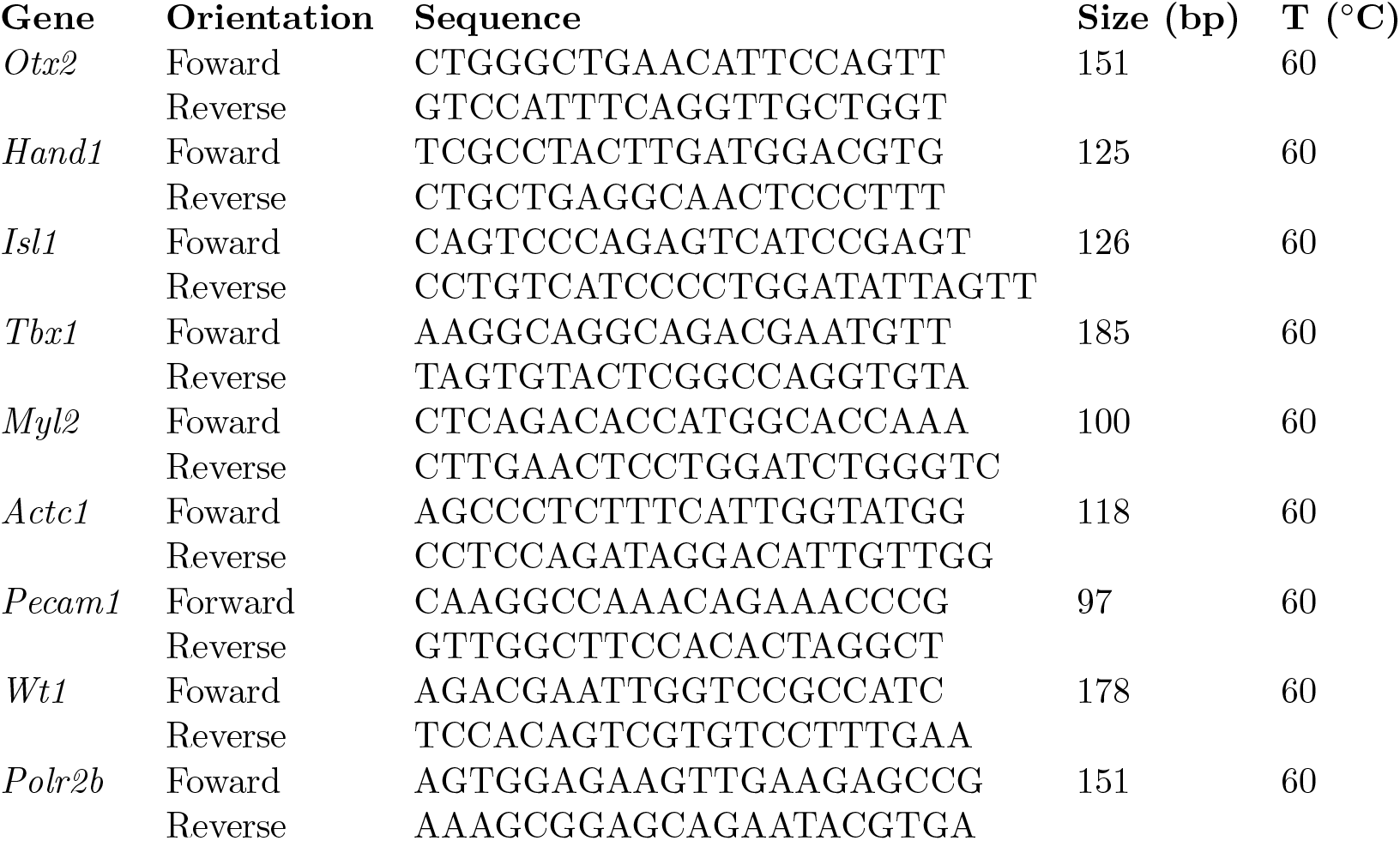
Primer sequences.

### 6.10 Statistical analysis

The experiments were conducted on at least three independent batches of cells. The measurements of data dispersion were calculated as standard deviation (SD). Comparisons between two experimental conditions were performed on the mean using a Student two-tailed test using Matlab (Mathworks). When more than two experimental groups were compared, an ANOVA was calculated with Holm–Bonferroni multi-comparison tests using Matlab (Mathworks). * = *p <* 0.05; ** = *p <* 0.01; * * * = *p <* 0.001.

## 7 Author Contributions

Conceptualization S.S., N.V.-Q. and C.N.B.; methodology, S.S. and N.V.-Q.; investigation and analysis S.S. and N.V.-Q.; image processing N.V.-Q., S.S. and A.A.; mESC pretreatment for gastruloid generation by T.T. and J.-C.C.; polarization analysis C.D.; writing – original draft, N.V.-Q. and S.S.; writing – review & editing, all authors; project supervision S.S.; funding acquisition C.N.B.

## 8 Acknowledgments

We acknowledge the Biomaterials and Microfluidics Core Facility (BMCF) and the FabLab (FLIP) of Institut Pasteur for the access to their microfabrication platforms. We thank Agnes Dubois from the Epigenomics, Proliferation and the Identity of Cells Lab at Institut Pasteur for providing the Rex1::GFP ESC reporter cell line, with the permission of Austin Smith from the University of Exeter. Sigolene Meilhac and Laurent Guillemot are acknowledged from the Heart Morphogenesis lab of Institut Pasteur for providing their expertise on the cardioid model.

## 10 Supplementary Information

**Supplementary Movie 1: Time evolution of mESCs aggregates in droplets**. 96 hours time-lapse microscopy of mESCs aggregating and proliferating at the bottom of 2 immobilized droplets in the microfluidic device. Scale bar represents 100 *μ*m.

**Supplementary Movie 2: A cardioid cultured in a well**. Real time microscopy movie of a beating D10 cardioid cultured in a well. Scale bar represents 100 *μ*m.

**Supplementary Movie 3: A cardioid cultured in the microfluidic device**. Real time microscopy movie of a beating D10 cardioid cultured in a droplet. Scale bar represents 100 *μ*m.

**Figure S1:**
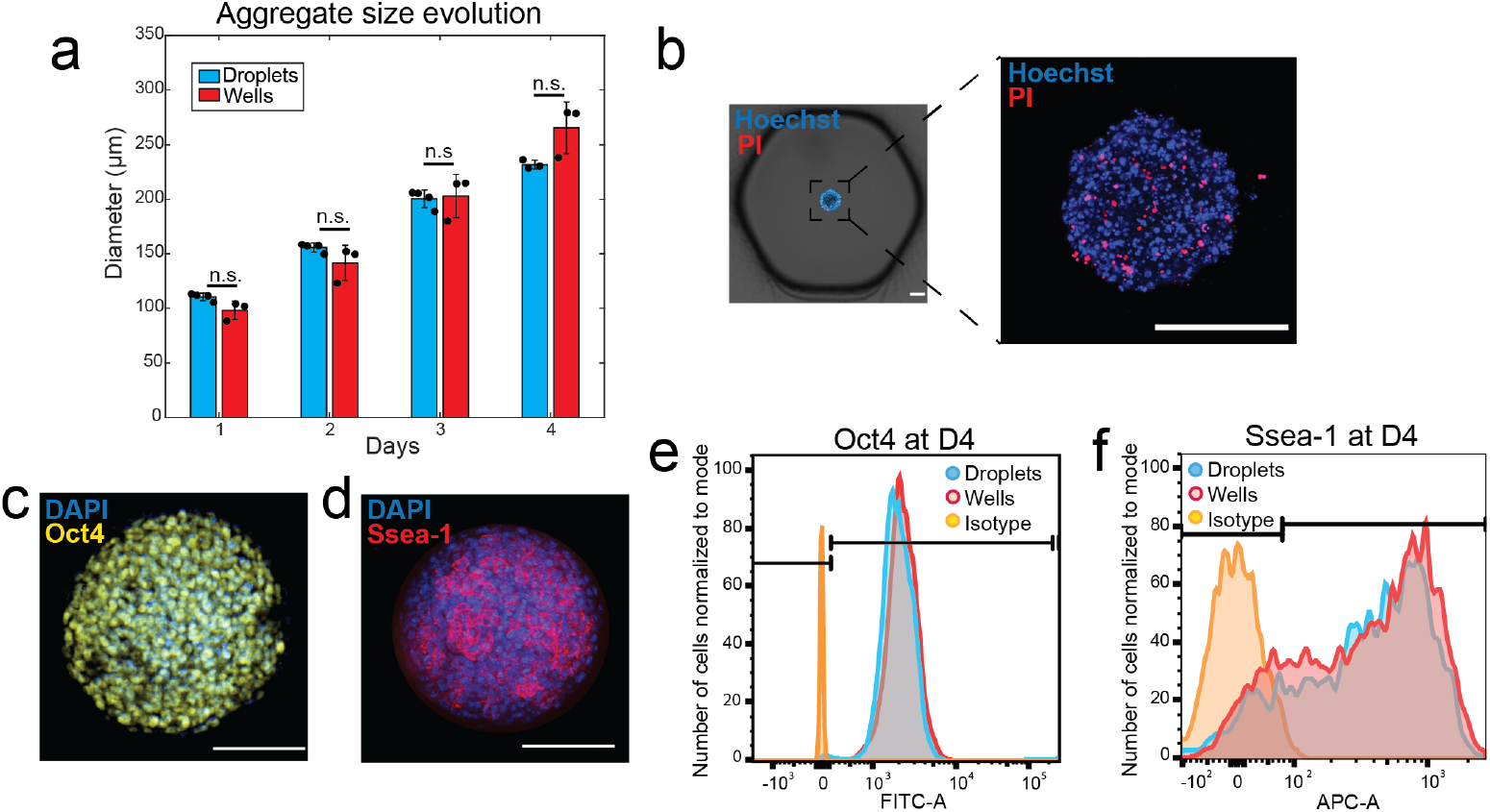
Culture of undifferentiated mESCs aggregates in microscale droplets. **(a)** Comparative of aggregate size evolution for aggregates cultured in 96-well plate *vs* chip (N=3 chips; N=3 plates). **(b)** Fluorescence image of a D4 aggregate stained with PI for viability assessment (out of three independent experiments). Scale bars represent 100 *μ*m. **(c-d)** Fluorescence images of an aggregate (out of three independent experiments) after 4 days of culture in the microfluidic device in 2iL conditions (CM^+^): in blue DAPI staining, in **(c)** yellow for Oct4 and in **(d)** Ssea-1 in red. Scale bars represent 100 *μ*m. **(e-f)** Flow cytometry plots of mESCs from aggregates cultured in chip vs 96-well plates, on **(h)** Oct4 and **(i)** Ssea-1.

**Figure S2:**
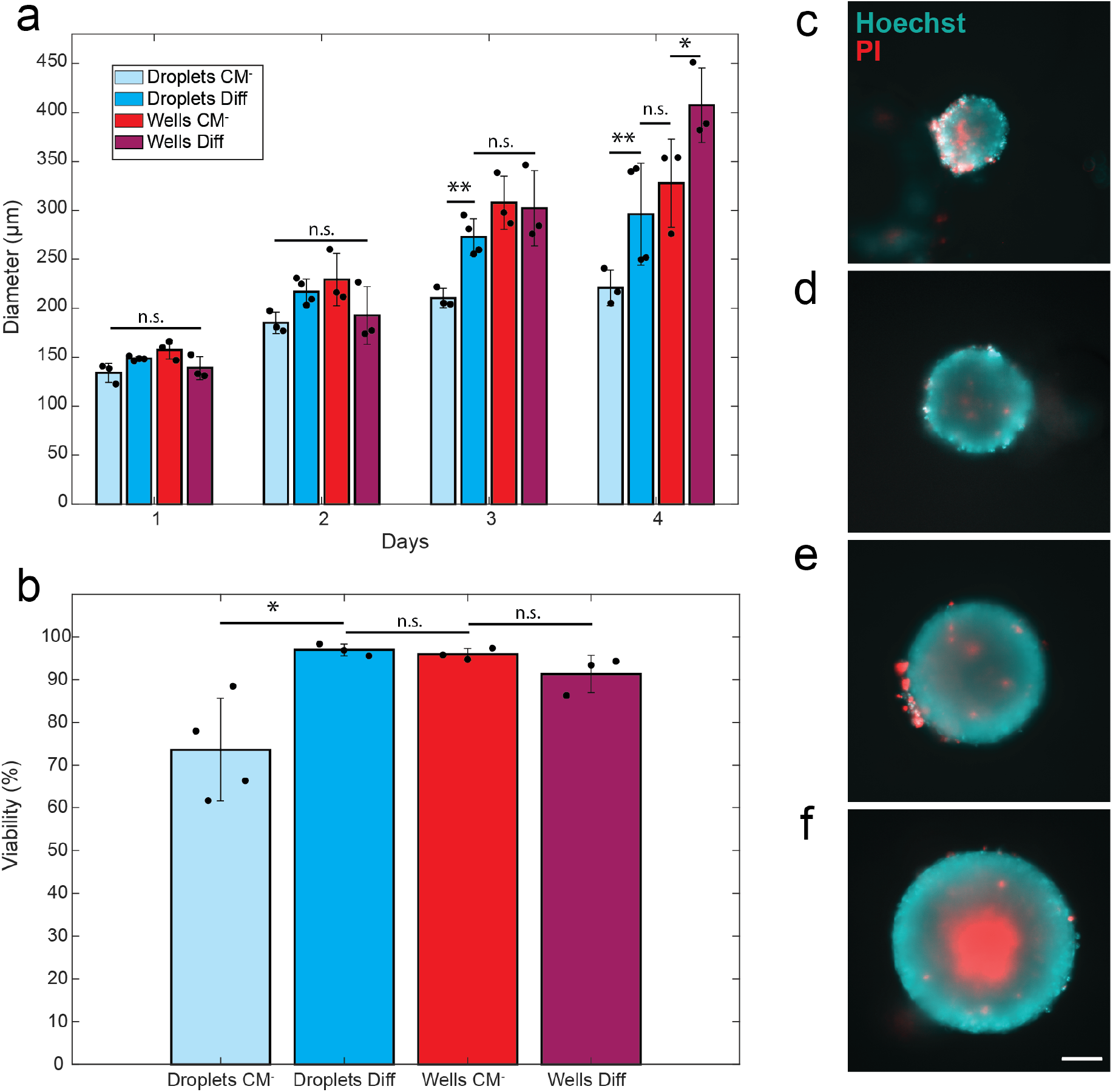
Screening of media for EB culture in droplets. **(a)** Evolution of aggregates diameter with different culture media over 4 days in culture (N=3 chips; N=3 plates). **(b)** Viability of cells in aggregates cultivated in different media at D4 (N=3 chips; N=3 plates). **(c)** Image of a cell aggregate cultivated in droplets (chip) in CM^-^. **(d)** Image of a D4 cell aggregate cultivated in wells in CM^-^. **(e)** Image of a D4 cell aggregate cultivated in droplets in Diff. **(f)** Image of a D4 cell aggregate cultivated in wells in Diff. Scale bar indicates 100 *μ*m.

**Figure S3:**
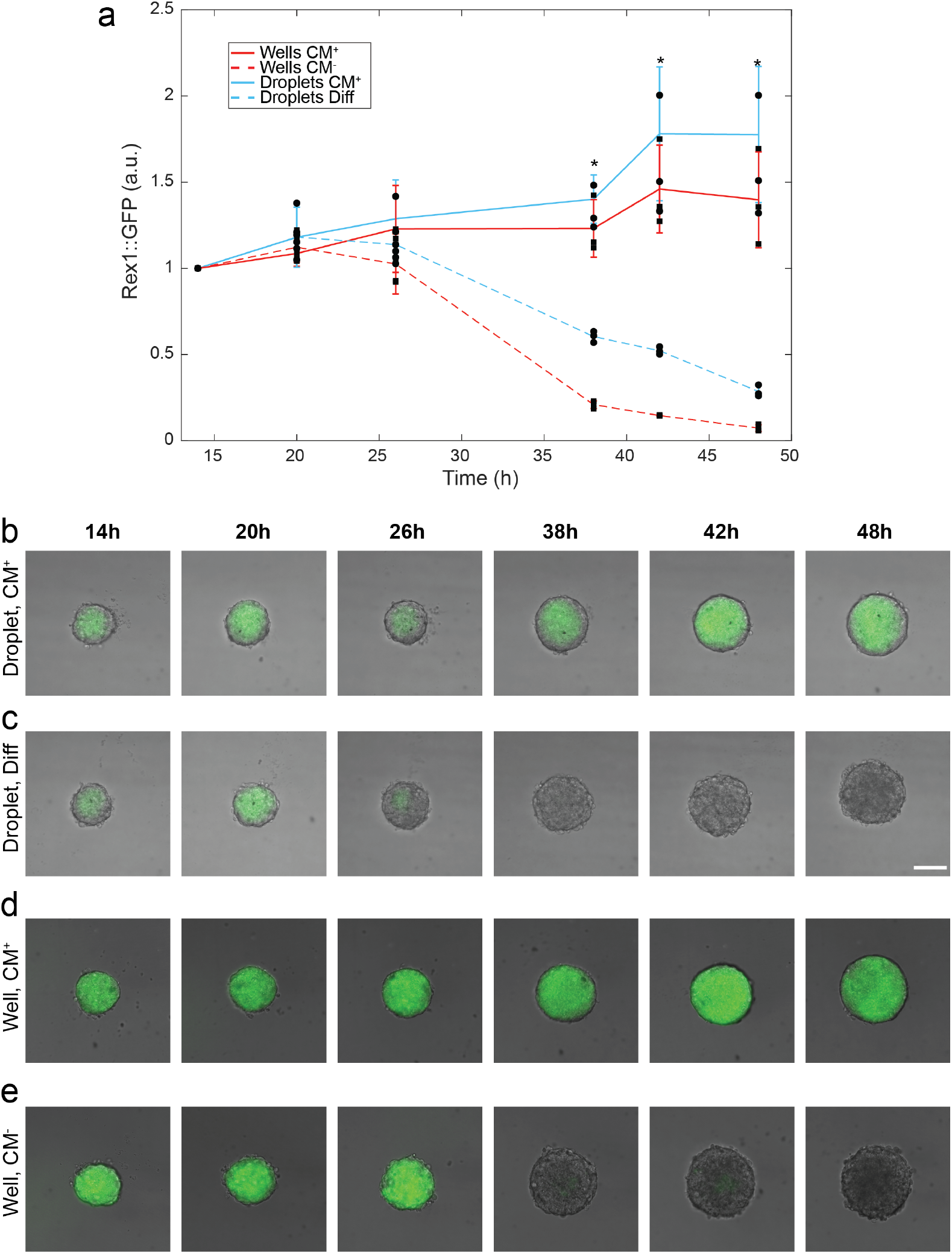
Rex1::GFP expression in mESCs aggregates cultivated in CM^-.^ over the first 48 hours of *in vitro* culture. **(a)** Dynamics of the expression of Rex1 (pluripotency marker) in aggregates cultured in droplets (CM^+^ or Diff) and in wells (CM^+^ or CM^-^)(N=3 chips; N=3 plates) CM^+^ was used as control in both conditions. Example of a mESCs aggregate cultured in **(b)** CM^+^ in a droplet, **(c)** Diff in a droplet, **(d)** CM^+^ in a well or **(e)** CM^-^ in a well. Scale bar represents 100 *μ*m.

**Figure S4:**
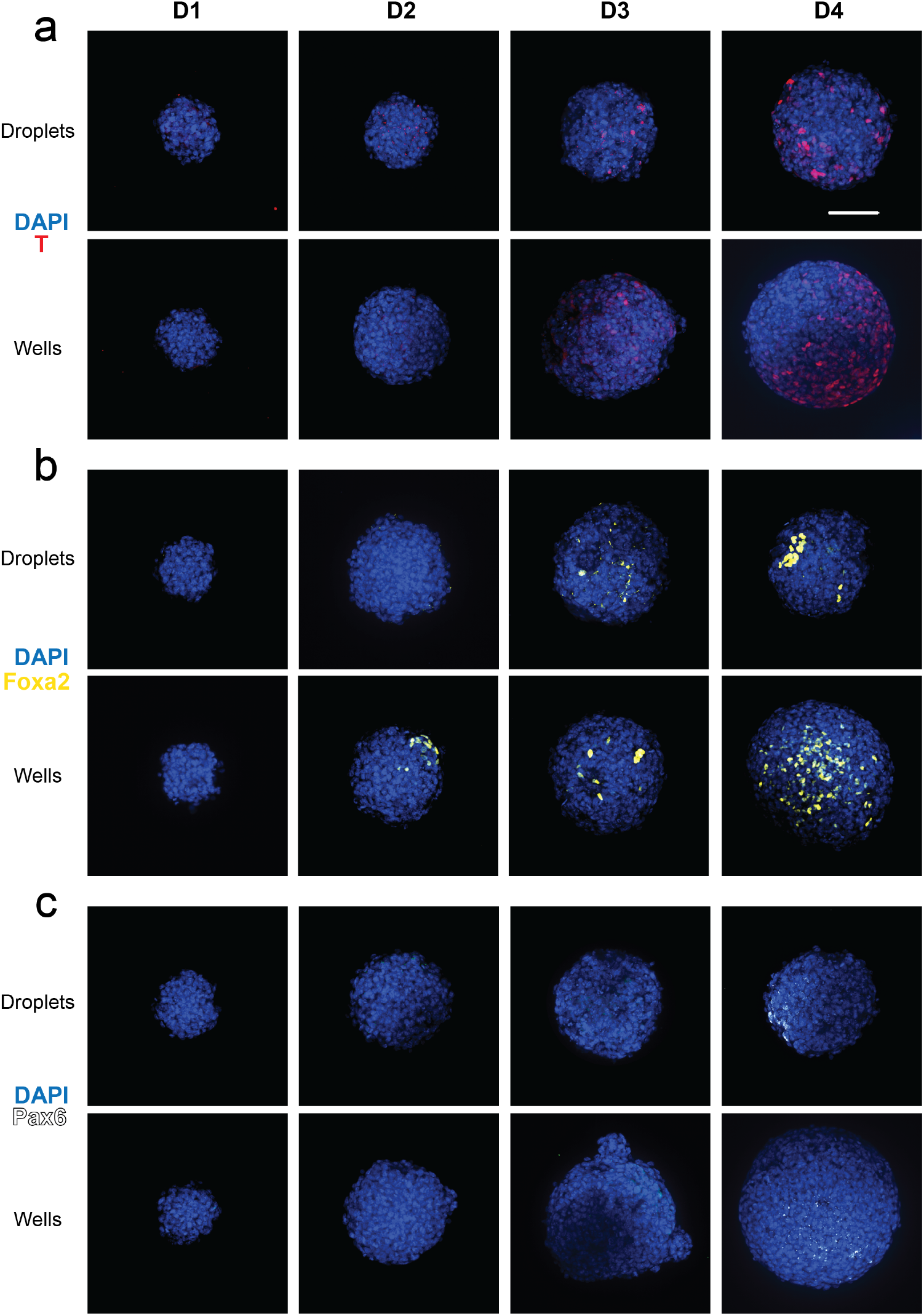
Cell differentiation on mESCs aggregates over the 4-days culture period in chips or in wells. Appearance of **(a)** Brachyury (T) positive cells (mesoderm marker), **(b)** Foxa2 positive cells (endoderm marker) and **(c)** Pax6 positive cells (ectoderm marker) in mESCs aggregates cultured for 4 days in chips and wells. Scale bar indicates 100 *μ*m.

**Figure S5:**
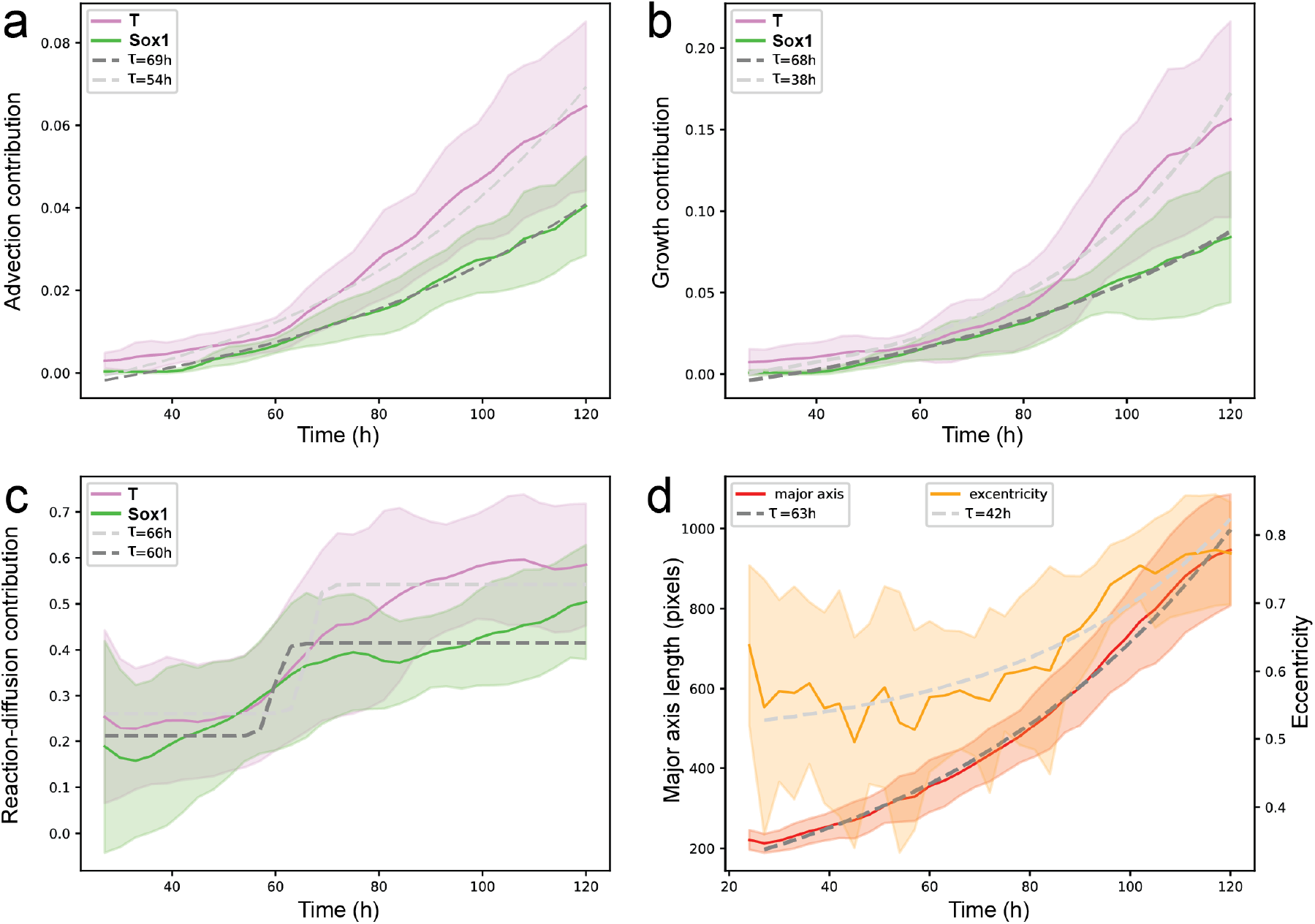
Best fit curve on the evolution of the contribution of advection, growth, reaction diffusion, major axis and eccentricity to gastruloid polarization. **(a)** Advection data (solid lines) and **(b)**growth data (solid lines) are fitted with exponential curves (doted lines). **(c)** reaction-diffusion data (solid lines) are fitted with logistic curves (doted lines). **(d)** The evolution of the gastruloid’s geometry is assessed via major axis length and eccentricity measurements, with both data (solid lines) fitted with exponential curves (dotted lines) (n = 9 gastruloids).

**Figure S6:**
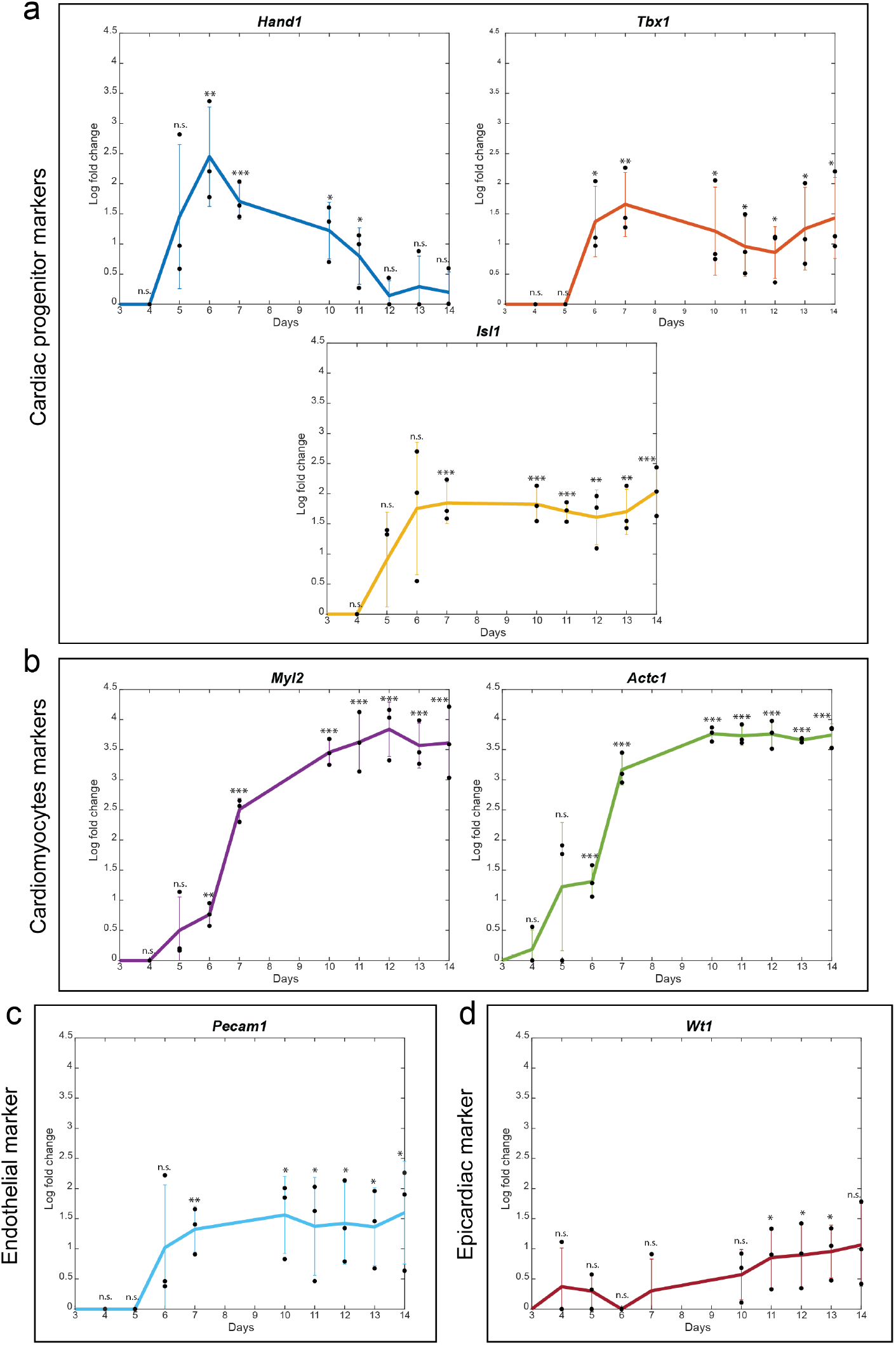
Evolution of the expression levels of cardiac differentiation markers over 14 days of culture. **(a)** Expression of cardiac progenitor markers: *Hand1, Tbx1* and *Isl1*. **(b)** Expression of cardiomyocytes markers: *Myl2* and *Actc1*. **(c)** Expression of the endothelial marker *Pecam1* **(d)** Expression of the epicardiac marker *Wt1* (N=3).

## References

[1] Sunghee Estelle Park, Andrei Georgescu, and Dongeun Huh. Organoids-on-a-chip. Science, 364(6444):960–965, 2019.

[2] Jeffrey M Karp, Judy Yeh, George Eng, Junji Fukuda, James Blumling, Kahp-Yang Suh, Jianjun Cheng, Alborz Mahdavi, Jeffrey Borenstein, Robert Langer, et al. Controlling size, shape and homogeneity of embryoid bodies using poly (ethylene glycol) microwells. Lab on a Chip, 7(6):786–794, 2007.

[3] EJ Vrij, S Espinoza, M Heilig, A Kolew, M Schneider, CA Van Blitterswijk, R. Truckenmüller, and NC Rivron. 3d high throughput screening and profiling of embryoid bodies in thermoformed microwell plates. Lab on a Chip, 16(4):734–742, 2016.

[4] Sébastien Sart, Gustave Ronteix, Shreyansh Jain, Gabriel Amselem, and Charles N Baroud. Cell culture in microfluidic droplets. Chemical Reviews, 122(7):7061–7096, 2022.

[5] Mohammad Mahfuz Chowdhury, Hiroshi Kimura, Teruo Fujii, and Yasuyuki Sakai. Induction of alternative fate other than default neuronal fate of embryonic stem cells in a membrane-based two-chambered microbioreactor by cell-secreted bmp4. Biomicrofluidics, 6(1):014117, 2012.

[6] Yusuke Sakai, Yukiko Yoshiura, and Kohji Nakazawa. Embryoid body culture of mouse embryonic stem cells using microwell and micropatterned chips. Journal of bioscience and bioengineering, 111(1):85–91, 2011.

[7] Joshua Guild, Amranul Haque, Pantea Gheibi, Yandong Gao, Kyung Jin Son, Elena Foster, Sophie Dumont, and Alexander Revzin. Embryonic stem cells cultured in microfluidic chambers take control of their fate by producing endogenous signals including lif. Stem Cells, 34(6):1501–1512, 2016.

[8] Laralynne Przybyla and Joel Voldman. Probing embryonic stem cell autocrine and paracrine signaling using microfluidics. Annual review of analytical chemistry, 5:293–315, 2012.

[9] Camilla Luni, Onelia Gagliano, and Nicola Elvassore. Derivation and differentiation of human pluripotent stem cells in microfluidic devices. Annual Review of Biomedical Engineering, 24:231–248, 2022.

[10] Makenzie G Bonner, Hemanth Gudapati, Xingrui Mou, and Samira Musah. Microfluidic systems for modeling human development. Development, 149(3):dev199463, 2022.

[11] Yaqing Wang, Hui Wang, Pengwei Deng, Wenwen Chen, Yaqiong Guo, Tingting Tao, and Jianhua Qin. In situ differentiation and generation of functional liver organoids from human ipscs in a 3d perfusable chip system. Lab on a Chip, 18(23):3606–3616, 2018.

[12] Yaqing Wang, Li Wang, Yaqiong Guo, Yujuan Zhu, and Jianhua Qin. Engineering stem cell-derived 3d brain organoids in a perfusable organ-on-a-chip system. RSC advances, 8(3):1677–1685, 2018.

[13] Fang Yu, Walter Hunziker, and Deepak Choudhury. Engineering microfluidic organoid-on-a-chip platforms. Micromachines, 10(3):165, 2019.

[14] Gloria Saorin, Isabella Caligiuri, and Flavio Rizzolio. Microfluidic organoids-on-a-chip: The future of human models. In Seminars in cell & developmental biology, volume 144, pages 41–54. Elsevier, 2023.

[15] Bas van Loo, Simone A Ten Den, Nuno Araújo-Gomes, Vincent de Jong, Rebecca R Snabel, Maik Schot, José M Rivera-Arbeláez, Gert Jan C Veenstra, Robert Passier, Tom Kamperman, et al. Mass production of lumenogenic human embryoid bodies and functional cardiospheres using in-air-generated microcapsules. Nature Communications, 14(1):6685, 2023.

[16] Adrien Saint-Sardos, Sebastien Sart, Kevin Lippera, Elodie Brient-Litzler, Sebastien Michelin, Gabriel Amselem, and Charles N Baroud. High-throughput measurements of intra-cellular and secreted cytokine from single spheroids using anchored microfluidic droplets. Small, 16(49):2002303, 2020.

[17] Jack-Christophe Cossec, Tatiana Traboulsi, Sébastien Sart, Yann Loe-Mie, Manuel Guthmann, Ivo A Hendriks, Ilan Theurillat, Michael L Nielsen, Maria-Elena Torres-Padilla, Charles N Baroud, et al. Transient suppression of sumoylation in embryonic stem cells generates embryo-like structures. Cell Reports, 42(4), 2023.

[18] Alexander Kumachev, Jesse Greener, Ethan Tumarkin, Erika Eiser, Peter W Zandstra, and Eugenia Kumacheva. High-throughput generation of hydrogel microbeads with varying elasticity for cell encapsulation. Biomaterials, 32(6):1477–1483, 2011.

[19] Hans Kleine-Brüggeney, Liisa D Van Vliet, Carla Mulas, Fabrice Gielen, Chibeza C Agley, José CR Silva, Austin Smith, Kevin Chalut, and Florian Hollfelder. Long-term perfusion culture of monoclonal embryonic stem cells in 3d hydrogel beads for continuous optical analysis of differentiation. Small, 15(5):1804576, 2019.

[20] Tina Tronser, Anna A Popova, Mona Jaggy, Martin Bastmeyer, and Pavel A Levkin. Droplet microarray based on patterned superhydrophobic surfaces prevents stem cell differentiation and enables high-throughput stem cell screening. Advanced healthcare materials, 6(23):1700622, 2017.

[21] Karen Lemke, Tobias Förster, Robert Römer, Mandy Quade, Stefan Wiedemeier, Andreas Grodrian, and Gunter Gastrock. A modular segmented-flow platform for 3d cell cultivation. Journal of Biotechnology, 205:59–69, 2015.

[22] Yanxi Liu, Shraddha Chakraborty, Chatrawee Direksilp, Johannes M Scheiger, Anna A Popova, and Pavel A Levkin. Miniaturized droplet microarray platform enables maintenance of human induced pluripotent stem cell pluripotency. Materials Today Bio, 12:100153, 2021.

[23] Yanxi Liu, Sarah Bertels, Markus Reischl, Ravindra Peravali, Martin Bastmeyer, Anna A Popova, and Pavel A Levkin. Droplet microarray based screening identifies proteins for maintaining pluripotency of hipscs. Advanced Healthcare Materials, 11(18):2200718, 2022.

[24] Huei-Wen Wu, Yi-Hsing Hsiao, Chih-Chen Chen, Shaw-Fang Yet, and Chia-Hsien Hsu. A pdms-based microfluidic hanging drop chip for embryoid body formation. Molecules, 21(7):882, 2016.

[25] Tatiana Traboulsi, Sébastien Sart, Charles N Baroud, Anne Dejean, and Jack-Christophe Cossec. Generation of embryo-like structures from mouse embryonic stem cells treated with a chemical inhibitor of sumoylation and cultured in microdroplets. STAR protocols, 4(4):102573, 2023.

[26] Sébastien Sart, Raphaël F-X Tomasi, Gabriel Amselem, and Charles N Baroud. Multiscale cytometry and regulation of 3d cell cultures on a chip. Nature communications, 8(1):469, 2017.

[27] Sébastien Sart, Raphaël F-X Tomasi, Antoine Barizien, Gabriel Amselem, Ana Cumano, and Charles N Baroud. Mapping the structure and biological functions within mesenchymal bodies using microfluidics. Science Advances, 6(10):eaaw7853, 2020.

[28] Raphaël F-X Tomasi, Sébastien Sart, Tiphaine Champetier, and Charles N Baroud. Individual control and quantification of 3d spheroids in a high-density microfluidic droplet array. Cell reports, 31(8):107670, 2020.

[29] Erwin Berthier, Ashley M Dostie, Ulri N Lee, Jean Berthier, and Ashleigh B Theberge. Open microfluidic capillary systems. Analytical chemistry, 91(14):8739–8750, 2019.

[30] Derk ten Berge, Wouter Koole, Christophe Fuerer, Matt Fish, Elif Eroglu, and Roel Nusse. Wnt signaling mediates self-organization and axis formation in embryoid bodies. Cell stem cell, 3(5):508–518, 2008.

[31] Macarena Lolas, Pablo DT Valenzuela, Robert Tjian, and Zhe Liu. Charting brachyury-mediated developmental pathways during early mouse embryogenesis. Proceedings of the National Academy of Sciences, 111(12):4478–4483, 2014.

[32] Antonio Simeone. Otx1 and otx2 in the development and evolution of the mammalian brain. The EMBO Journal, 1998.

[33] Susanne C Van den Brink, Peter Baillie-Johnson, Tina Balayo, Anna-Katerina Hadjantonakis, Sonja Nowotschin, David A Turner, and Alfonso Martinez Arias. Symmetry breaking, germ layer specification and axial organisation in aggregates of mouse embryonic stem cells. Development, 141(22):4231–4242, 2014.

[34] David A Turner, Mehmet Girgin, Luz Alonso-Crisostomo, Vikas Trivedi, Peter Baillie-Johnson, Cherise R Glodowski, Penelope C Hayward, Jérôme Collignon, Carsten Gustavsen, Palle Serup, et al. Anteroposterior polarity and elongation in the absence of extra-embryonic tissues and of spatially localised signalling in gastruloids: mammalian embryonic organoids. Development, 144(21):3894–3906, 2017.

[35] Simon Gsell, Sham Tlili, Matthias Merkel, and Pierre-François Lenne. Marangoni-like tissue flows enhance symmetry breaking of embryonic organoids. bioRxiv, pages 2023–09, 2023.

[36] Ségoléne Bernheim, Adrien Borgel, Jean-François Le Garrec, Emeline Perthame, Audrey Desgrange, Cindy Michel, Laurent Guillemot, Sébastien Sart, Charles N. Baroud, Wojciech Krezel, Francesca Raimondi, Damien Bonnet, Stéphane Zaffran, Lucile Houyel, and Sigoléne M. Meilhac. Torsion of the heart tube by shortage of progenitor cells : identification of greb1l as a genetic determinant of criss-cross heart in mice. Developmental Cell, 2023.

[37] Nan Cao, Zumei Liu, Zhongyan Chen, Jia Wang, Taotao Chen, Xiaoyang Zhao, Yu Ma, Lianju Qin, Jiuhong Kang, Bin Wei, et al. Ascorbic acid enhances the cardiac differentiation of induced pluripotent stem cells through promoting the proliferation of cardiac progenitor cells. Cell research, 22(1):219–236, 2012.

[38] Deepti Abbey and Polani B Seshagiri. Ascorbic acid-mediated enhanced cardiomyocyte differentiation of mouse es-cells involves interplay of dna methylation and multiple-signals. Differentiation, 96:1–14, 2017.

[39] Whitney Edwards, Todd M Greco, Gregory E Miner, Natalie K Barker, Laura Herring, Sarah Cohen, Ileana M Cristea, and Frank L Conlon. Quantitative proteomic profiling identifies global protein network dynamics in murine embryonic heart development. Developmental Cell, 2023.

[40] Richard CV Tyser, Ximena Ibarra-Soria, Katie McDole, Satish Arcot Jayaram, Jonathan Godwin, Teun AH van den Brand, Antonio MA Miranda, Antonio Scialdone, Philipp J Keller, John C Marioni, et al. Characterization of a common progenitor pool of the epicardium and myocardium. Science, 371(6533):eabb2986, 2021.

[41] Kejie Chen, Yi Zheng, Xufeng Xue, Yue Liu, Agnes M Resto Irizarry, Huaijing Tang, and Jianping Fu. Branching development of early post-implantation human embryonic-like tissues in 3d stem cell culture. Biomaterials, 275:120898, 2021.

[42] Giovanni G Giobbe, Monica Zagallo, Massimo Riello, Elena Serena, Giulia Masi, Luisa Barzon, Barbara Di Camillo, and Nicola Elvassore. Confined 3d microenvironment regulates early differentiation in human pluripotent stem cells. Biotechnology and bioengineering, 109(12):3119–3132, 2012.

[43] Jiani Cao, Meng Li, Kun Liu, Xingxing Shi, Ning Sui, Yuchen Yao, Xiaojing Wang, Shaojing Tan, Qian Zhao, Liang Wang, et al. Oxidative phosphorylation safeguards pluripotency via udp-n-acetylglucosamine. bioRxiv, pages 2021–07, 2021.

[44] Tüzer Kalkan, Nelly Olova, Mila Roode, Carla Mulas, Heather J Lee, Isabelle Nett, Hendrik Marks, Rachael Walker, Hendrik G Stunnenberg, Kathryn S Lilley, et al. Tracking the embryonic stem cell transition from ground state pluripotency. Development, 144(7):1221–1234, 2017.

[45] David A Turner, Pau Rué, Jonathan P Mackenzie, Eleanor Davies, and Alfonso Martinez Arias. Brachyury cooperates with wnt/β-catenin signalling to elicit primitive-streak-like behaviour in differentiating mouse embryonic stem cells. BMC biology, 12:1–19, 2014.

[46] Steven A Jackson, Jacqueline Schiesser, Edouard G Stanley, and Andrew G Elefanty. Differentiating embryonic stem cells pass through ‘temporal windows’ that mark responsiveness to exogenous and paracrine mesendoderm inducing signals. Plos one, 5(5):e10706, 2010.

[47] Alyssa V Ngangan, James C Waring, Marissa T Cooke, Christian J Mandrycky, and Todd C McDevitt. Soluble factors secreted by differentiating embryonic stem cells stimulate exogenous cell proliferation and migration. Stem Cell Research & Therapy, 5:1–12, 2014.

[48] Tamar Dvash, Nadav Sharon, Ofra Yanuka, and Nissim Benvenisty. Molecular analysis of lefty-expressing cells in early human embryoid bodies. Stem Cells, 25(2):465–472, 2007.

[49] Berna Sozen Kaya, Gianluca Amadei, Andy Cox, Ran Wang, Ellen Na, Sylwia Czukiewska, Lia Chappell, Thierry Voet, Geert Michel, Naihe Jing, et al. Self-assembly of embryonic and two extra-embryonic stem cell types into gastrulating embryo-like structures. 2018.

[50] Guillaume Blin, Darren Wisniewski, Catherine Picart, Manuel Thery, Michel Puceat, and Sally Lowell. Geometrical confinement controls the asymmetric patterning of brachyury in cultures of pluripotent cells. Development, 145(18):dev166025, 2018.

[51] Naor Sagy, Shaked Slovin, Maya Allalouf, Maayan Pour, Gaya Savyon, Jonathan Boxman, and Iftach Nachman. Prediction and control of symmetry breaking in embryoid bodies by environment and signal integration. Development, 146(20):dev181917, 2019.

[52] Ali Hashmi, Sham Tlili, Pierre Perrin, Molly Lowndes, Hanna Peradziryi, Joshua M Brickman, Alfonso Martínez Arias, and Pierre-François Lenne. Cell-state transitions and collective cell movement generate an endoderm-like region in gastruloids. Elife, 11:e59371, 2022.

[53] Olivier Frey, Patrick M Misun, David A Fluri, Jan G Hengstler, and Andreas Hierlemann. Reconfigurable microfluidic hanging drop network for multi-tissue interaction and analysis. Nature communications, 5(1):4250, 2014.

[54] Yonatan R Lewis-Israeli, Aaron H Wasserman, Mitchell A Gabalski, Brett D Volmert, Yixuan Ming, Kristen A Ball, Weiyang Yang, Jinyun Zou, Guangming Ni, Natalia Pajares, et al. Self-assembling human heart organoids for the modeling of cardiac development and congenital heart disease. Nature communications, 12(1):5142, 2021.

[55] Pablo Hofbauer, Stefan M Jahnel, Nora Papai, Magdalena Giesshammer, Alison Deyett, Clara Schmidt, Mirjam Penc, Katherina Tavernini, Nastasja Grdseloff, Christy Meledeth, et al. Cardioids reveal self-organizing principles of human cardiogenesis. Cell, 184(12):3299–3317, 2021.

[56] Cédric Deluz, Elias T Friman, Daniel Strebinger, Alexander Benke, Mahé Raccaud Andrea Callegari, Marion Leleu, Suliana Manley, and David M Suter. A role for mitotic bookmarking of sox2 in pluripotency and differentiation. Genes & development, 30(22):2538–2550, 2016.

